# Necrosis triggered by *Mycobacterium tuberculosis* alters macrophage triglyceride metabolism and inflammatory response in a DGAT1 – dependent manner

**DOI:** 10.1101/187104

**Authors:** Neetika Jaisinghani, Stanzin Dawa, Kaurab Singh, Ananya Nandy, Dilip Menon, Purva Bhandari, Garima Khare, Anil Tyagi, Sheetal Gandotra

**Author notes:** **Materials and correspondence:** Sheetal Gandotra.

## Abstract

Tuberculosis granulomas represent a site of controlling bacterial dissemination at the cost of host tissue damage. Within the heterogenous array of TB granulomas, some contain triglyceride (TG) rich foamy macrophages, the etiology and function of which remains largely unexplained. Here we show that necrosis of tuberculosis lesions and *M. tuberculosis* (Mtb) infected macrophages elicits a bystander response of triglyceride storage. We elucidate a role for the RD1 region of mycobacterial genome to be a key player in this phenomenon. TG storage in necrosis associated foamy macrophages promoted the pro-inflammatory state of macrophages while silencing expression of diacylglycerol O-acyltransferase (DGAT1) suppressed expression of pro-inflammatory genes. Surprisingly, acute infection with Mtb led to lipolysis of host TG, rather than synthesis, suggesting mobilization of triglyceride stored within macrophage lipid droplets to be involved in the inflammatory response to infection. Our data is likely to open new avenues for management of the inflammatory response during infection.

## Introduction

Tuberculosis is the major worldwide cause of mortality due to an infectious agent ^1^. The causative pathogen, *Mycobacterium tuberculosis*, evades host immunity and adapts to the host’s lipid rich environment ^23^. This lipid rich environment histologically presents as caseous necrosis and has been characterized biochemically to be a triglyceride rich ^4^. The transformation of a nascent granuloma to this state is not fully understood and the contribution of triglycerides in host immunity remains elusive.

The importance of lipid metabolism in infection induced inflammatory response has emerged from benefits of treatments that target lipid metabolism in pre-clinical models of tuberculosis. These models have revealed that nuclear receptors responsive to lipids, such as peroxisome proliferator-activated receptor-γ (PPAR*γ*) and liver X receptors (LXRα and LXRβ) protect against tuberculosis by regulating both pro and anti-inflammatory pathways ^56^. The absence of LXRα and LXRβ led to increased bacterial burden and lowering of the protective early neutrophil recruitment and Th17 response. In addition, treatment of infected mice with an agonist of LXR led to control in bacterial numbers and increase in the Th17 response. The antimicrobial effect is thought to be largely through LXR dependent inhibition of cholesterol efflux, which limits cholesterol availability for mycobacteria^7^. Along the same lines, targeting the synthesis of lipid mediators of inflammation derived from arachidonic acid, by the non-steroidal anti-inflammatory drug Ibuprofen, has shown promise towards limiting pathology in the mouse model of infection 8. Given that triglyceride is a major lipid in human TB granulomas and acts as a sink of fatty acids and signalling mediators such as diacylglycerides, phosphatidic acid, and phospholipids, its role in TB infection needs to be addressed. Understanding the mechanistic basis for differentiation of macrophages to triglyceride rich foamy macrophages during infection will guide research efforts towards overcoming inflammation associated with morbidity due to tuberculosis.

Several models have been described for increased lipid biogenesis in macrophages upon *M. tuberculosis* infection. As mycobacterial surface is enriched in non-polar lipids, these lipids are proposed to promote the emergence of neutral lipid rich macrophages ^9,10^. Alternative models of increased *de novo* fatty acid synthesis and inhibition of lipolysis mediated by perilipin 1A have also been suggested ^11^. However, in human tuberculosis, neutral lipid rich macrophages are conspicuously absent in solid granulomas and present in necrotic granulomas ^10^. Therefore, the direct role of mycobacterial lipids or active rewiring of macrophage metabolism by intracellular mycobacteria may be insufficient to explain the emergence of foamy macrophages *in vivo*.

The role of the host immune response in emergence of triglyceride rich foamy macrophages has been suggested by others. For example, KN Balaji’s group found that TLR2 deletion leads to reduction in the ability of infected mice to develop foamy macrophages ^12^. It is important to note that macrophage TLR2 expression is required for induction of pro-inflammatory cytokines in response to Mtb infection ^13^ while TLR2^-/^- mice exhibit higher expression of TNFα and IL12p40 during chronic infection ^14^. Therefore the role of inflammation in driving the emergence of foamy macrophages *in vivo* cannot be ruled out. In addition, LXRα and LXRβ, regulators of inflammation during murine tuberculosis, are suggested to be negative regulators of foamy macrophages due to higher neutral lipid accumulation in lesions from LXRαβ^-/-^ mice compared to wild type C57BL/6 mice ^6^. Both TLR2^-/^- and LXR^-/^- mice exhibit heightened pro-inflammatory responses upon TB infection with aggravated pathology involving necrosis. Concomitant to the dual role played by the local inflammatory response are the architectural changes that provide heterogeneity amongst TB lesions within the same genetic background ^15,16^. Therefore, a role of the incident pathology in differentiation of macrophages to foamy macrophages needs to be addressed.

Death of infected macrophages as a host response during tuberculosis decides the fate for bacterial control versus dissemination where apoptosis contains bacteria via efferocytosis and necrosis helps in dissemination of Mtb ^17,18^. Virulent strains of Mtb are known to inhibit apoptosis and induce necrosis in infected macrophages as a mechanism to evade their own death ^19-21^.The inflammatory response in TB also contributes to tissue necrosis and cavitation which is the characteristic end-stage tissue damage ^22,23^ leading to chronic impairment of pulmonary function ^24,25^. Using macrophage and mouse models of infection, the RD1 locus of *M. tuberculosis* has been shown to contribute to necrosis induced upon Mtb infection ^26,27^. This locus encodes the mycobacterial pathogenesis associated ESX-1 secretion system, which is responsible for damage to the host membrane ^28,29^. In an attempt to understand the kinetics and underlying mechanism of foamy macrophage formation during Mtb infection, we discovered a role for RD1 mediated necrosis in the differentiation of macrophages towards triglyceride-rich, pro-inflammatory foamy macrophages. We studied the appearance of foamy macrophages at different stages in the guinea pig model of tuberculosis infection and in Mtb infected THP1 and primary human macrophages, leading to the development of an *in vitro* model of necrosis-associated foamy macrophages. We elucidate exogenous FA to be the carbon source for TG biosynthesis in response to necrosis. These necrosis associated foamy macrophages display a hyper-inflammatory response to infection *in vitro* and *in vivo*. The pro-inflammatory nature of these foamy macrophages could be controlled by genetic silencing of the enzyme diacylglycerol O-acyltransferase (DGAT1). We demonstrate that TG biosynthesis in human macrophages is important for the innate immune response of macrophages to Mtb and may play a critical role in the ensuing inflammation during TB.

## Results

### Necrosis associated TG accumulation in guinea pig granulomas

The guinea pig model of pulmonary tuberculosis presents with the array of granulomas observed in human tuberculosis, including solid and necrotic granulomas which may be caseous, fibrocaseous, or cavitary lesions ^4,30,31^. We investigated the presence of TG rich foamy macrophages in solid and necrotic granulomas of guinea pig lungs post aerosol infection with a low dose of Mtb H37Rv. 10 sections from lungs of each animal at week 4 and week 10 post infection were analyzed. Out of 30 sections obtained from each time point, we found 27 granulomas at week 4 and 33 granulomas at week 10 post infection (Table 1). Only 2 granulomas at week 4 were necrotic (7.4%) as compared to 17 at week 10 (51.5%) post infection (Figure 1a). CFUs increased four-fold between week 4 and 10 (Figure 1b), consistent with association of necrosis with higher bacterial burden in the guinea pig model of tuberculosis ^32^. Staining of these granulomas revealed that 100% of necrotic granulomas (Table 1) were Oil red O positive. Oil red O positive macrophages (hereafter referred as foamy macrophages) were conspicuous around the necrotic core of the granuloma (Figure 1c and d). In contrast, solid granulomas were found to be devoid of any Oil red O staining (Table 1). One of the animals at week 10 was found to contain mostly Oil red O positive granulomas; lipid analysis from macroscopic granulomas resected from this animal verified an increase in TG content as compared to anatomically similar regions from an animal at week 4 post infection or an uninfected guinea pig (Figure 1e and Figure S1). These data demonstrated that as granulomas evolved to develop necrosis, they exhibited presence of TG rich foamy macrophages proximal to the necrotic core.

**Figure 1:**
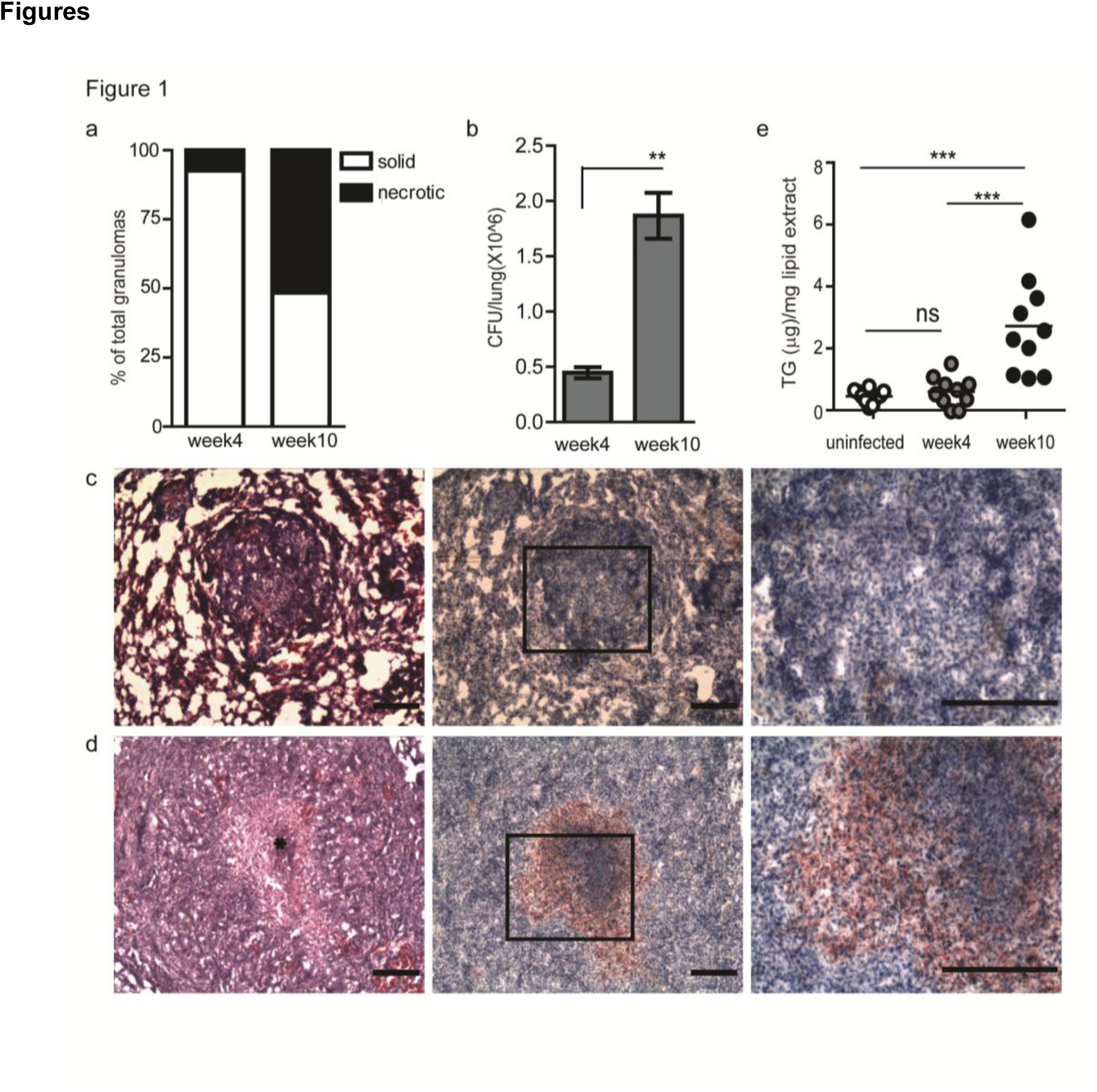
Necrotic granulomas contain triglyceride rich foamy macrophages while solid granulomas do not. (a) Percentage of solid and necrotic granulomas of total granulomas from tissue sections analysed from three animals at each time point. (b) CFU/ lung as calculated from left caudal lobe per animal from a total of three animals at each time point. Sequential cryosections stained with haematoxylin+eosin (H & E) (left panel) and haematoxylin+Oil red O (center and right panel) from week 4 (c) and 10 (d) infected guinea pig lungs. * indicates necrotic center filled with pyknotic nuclei. Scale bar=100 μm. (e) TG quantification from 10 individual granuloma sections from a single animal at each time point, as calculated using densitometry. Each circle represents TG from a single section. Line indicates mean TG of all 10 sections (**p<0.01, ***- p≤0.001, ns-not significant). See also Figure S1.

**Table 1:** Distribution of Oil red O positive cells (ORO^+^) across granuloma types

|  | ORO<sup>+</sup> solid granulomas | ORO<sup>+</sup> necrotic granulomas |
| --- | --- | --- |
| Week 4 (n=27) | 0/25 | 2/2 |
| Week 10 (n=33) | 0/16 | 17/17 |

### Necrosis and TG storage in Mtb infected macrophages

To test if necrosis of Mtb infected macrophages triggers the appearance of triglyceride rich foamy macrophages, we infected human macrophages with increasing multiplicity of infection (MOI) so as to induce necrosis as reported earlier ^19,33^. After infecting phorbol 12-myristate 13-acetate (PMA) differentiated THP1 macrophages with the virulent strain H37Rv for 24h, we observed 20% necrosis at MOI1 which increased to 50% at MOI10 and 70% at MOI50 (Figure 2a and Figure S3). To assess the effect of increased MOI on macrophage lipid metabolism we pulsed the macrophages with ^14^C-oleate to measure new triglyceride synthesis during infection. The addition of orlistat (10 μM) in the assay prevented lipolysis of newly synthesized TG and therefore allowed measurement of synthesis accurately ^34^ (Figure S2a). This concentration of orlistat did not impact the viability of Mtb within macrophages (Figure S2b). We found no difference in TG synthesis between uninfected and MOI1 infected cells (Figure 2b). Increasing MOI to 5 or 10 increased TG synthesis two folds and MOI 50 increased TG synthesis by eight folds (Figure 2b). TG and other neutral lipids are bound by a monolayer of phospholipid membrane within cytosolic lipid droplets (LDs)^35^. To understand how increasing MOI alters the total neutral lipid content in macrophages, we also measured the size of LDs in uninfected and Mtb infected macrophages using a neutral lipid stain BODIPY493/503 (Figure 2c). BODIPY493/503 was chosen over other commercially available neutral lipid stains such as LipidTox after ascertaining that the former did not stain bacterial surface both *in vitro* and *in vivo* as opposed to LipidTox, which could stain Mtb in addition to host LDs (Figure S4a, b and c). Mtb infection at MOI1^24h^, did not induce an increase in size of LDs, LD content per cell or total triglyceride content of macrophages (Figure 2c-e and Figure S4d, e). Human peripheral blood monocyte derived macrophages, similar to THP1 macrophages, contained a basal level of ADRP coated lipid droplets which did not increase in size or content upon infection at MOI1^24h^ (Figure S4f, g, and h). Increase in MOI to 10 and 50 led to an increase in the median size of the BODIPY493/503 stained LDs of THP1 macrophages (Figure 2c and d), consistent with the increase in FA assimilation into TG (Figure 2b). Furthermore, the increased FA assimilation into macrophage TG corresponded to an overall increase in TG content (Figure 2e). TG content in the remaining adherent cells revealed a linear correlation with the relative necrosis in that well (R^2^=0.59, p=0.0002) (Figure 2f). We further tested if fatty acids released from necrotic cells could contribute to the triglyceride pool of uninfected or bystander macrophages using a transwell assay. BODIPY558/568-C12 FA labelled lipids from MOI50^24h^ infected cells could be assimilated into the neutral lipid pool of uninfected cells across a 0.45|a filter (Figure 2g), confirming that lipids of cells undergoing necrosis upon Mtb infection can provide a source of fatty acids for triglyceride synthesis in bystander cells.

**Figure 2:**
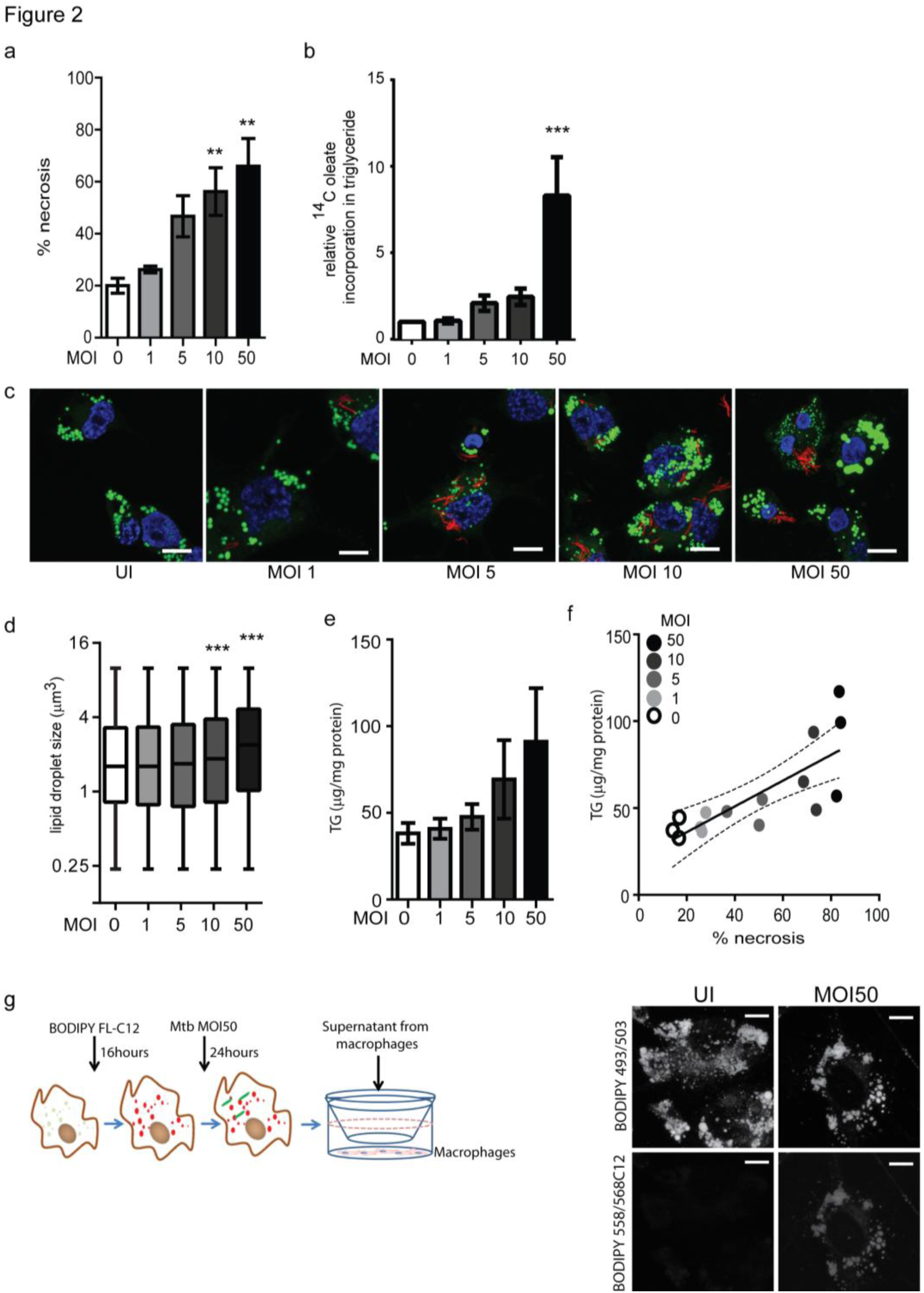
Increased TG synthesis during Mtb infection correlates with necrosis. (a) Quantification of cell death measured by LDH activity and represented as % of necrosis relative to detergent lysed cells. Data are mean ±SEM from 3 experiments. (b) Incorporation of ^14^C oleate into TG in Mtb infected macrophages at indicated multiplicities of infection (MOI) relative to uninfected macrophages. Data are mean ± SEM from 4 independent experiments. (c) BODIPY 493/503 (green) and DAPI (blue) stained THP1 macrophages at 24h post infection with indicated MOI. mCherry^+^ Mtb can be seen in red. Scale bar=10μM. (d) Assessment of individual LD size from 5 confocal z-stacks of each condition. Asterics indicate comparison of medians from 5,800-11,000 lipid droplets per group. (e) TG estimation in adherent cells at 24h post infection with indicated MOI. Data are mean ±SD from 3 wells. (f) Correlation of cell death in a well to the TG content of the remaining cells in that well. Each circle represents value from single well. Solid line represents line of linear regression of LDH released with TG content. Dotted lines represent confidence intervals of the best fit line (solid). (R^2^=0.59, p=0.0002) (g) (Left panel) Schematic representing experimental set-up for supernatant transfer from BODIPY558/568C12 labelled macrophages. (Right panel) BODIPY 493/503 staining in cells treated with supernatant from uninfected or *M. tuberculosis* infected macrophages labelled with BODIPY558/568C12 prior to infection. Scale bar=10μm. (**p<0.01, ***p<0.001). See also Figure S2, S3, and S4.

The mycobacterial pathogenicity associated locus, *region of difference 1* (RD1), has been shown to be involved in inducing atypical cellular necrosis ^19,27,36^ via a bacterial contact-dependent membrane deformation ^28^. The vaccine strain *M. bovis* BCG harbours a deletion of the RD1 locus ^37^ and is known to induce apoptosis rather than necrosis in macrophages ^38,39^. Consistent with literature, we also observed Mtb infection to cause necrosis instead of apoptosis with poor Annexin V staining (1-15%, N=3) and efficient propidium iodide (PI) staining (20-85%, N=3) (Figure 3a-c). In comparison, infection with ARD1 strain and *M. bovis* BCG led to 10-20% annexin V positivity and 10% or lower PI positivity (Figure 3a-c). LDH release assays confirmed that infection with ΔRD1 strain or *M. bovis* BCG induced significantly lower necrosis compared to wild type H37Rv at MOI5, 10 and 50 (Figure 3d, f and Figure S5a, b). The extent of MOI dependent increase in FA assimilation in case of H37Rv was not observed in case of the ΔRD1 strain (Figure 3e and g). Cell death upon infection at MOI10^24h^ of H37Rv and MOI50^24h^ of ΔRD1 was comparable (approximately 40%) (Figure 3d). In both cases, triglyceride synthesis from exogenous fatty acids was increased approximately 4-fold relative to uninfected cells (Figure 3e). To rule out the possibility of RD1 in regulating fatty acid uptake in macrophages independent of necrosis, we used a pharmacological inhibitor of necrosis to test our hypothesis. We found IM-54, an inhibitor of H_2_O_2_ induced necrosis ^40^, to inhibit necrosis in Mtb infected macrophages in a dose dependent manner (Figure S5c and Figure 3h). 20|aM IM-54 reduced cell death from 70% to 40% at MOI50^24h^. This concentration of IM-54 led to almost 50% reduction in ^14^C-oleate assimilation into TG (Figure 3i), confirming a role for infection induced necrosis to be a player in stimulating FA assimilation in bystander macrophages.

**Figure 3:**
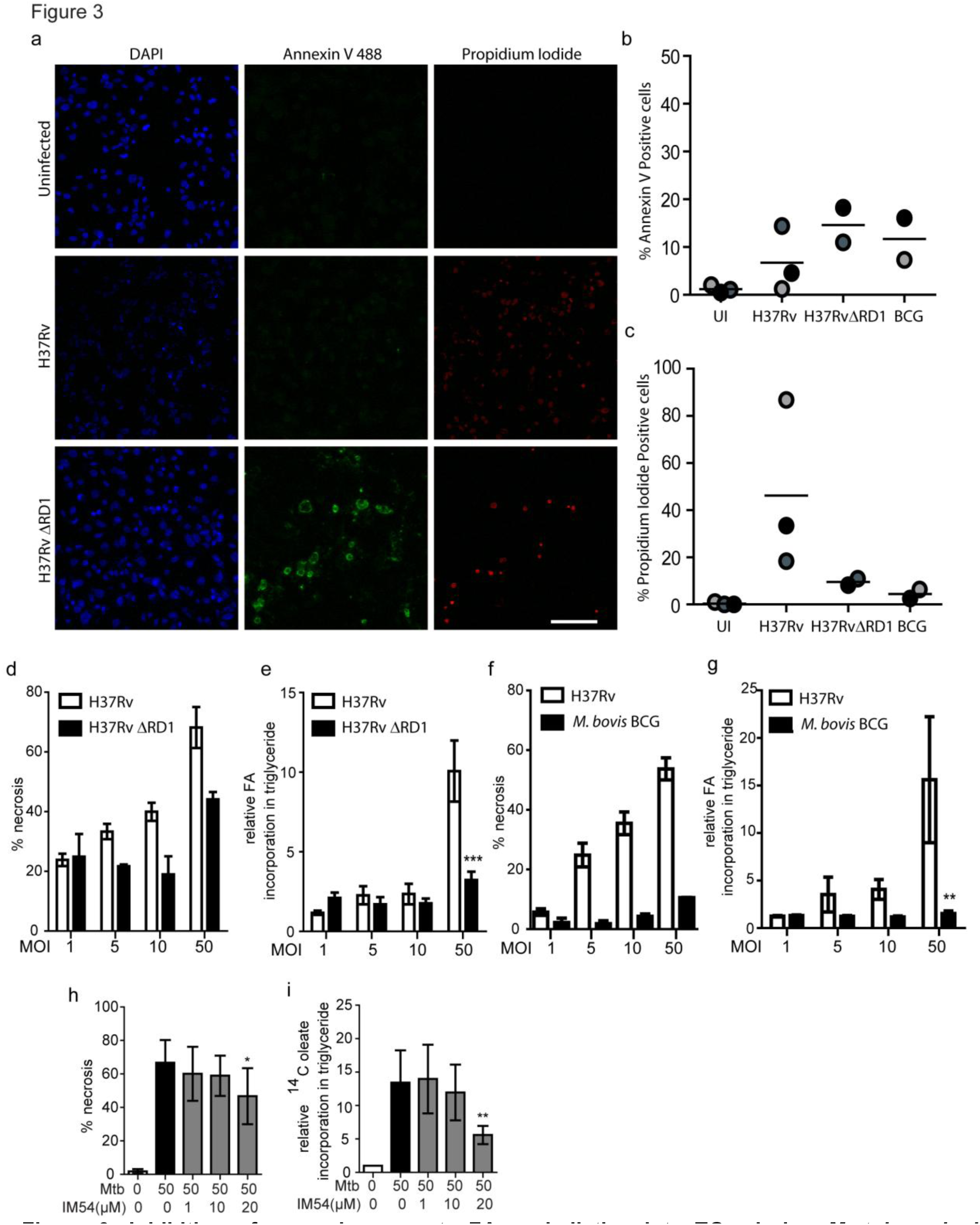
Inhibition of necrosis prevents FA assimilation into TGs during *M. tuberculosis* infection. (a) Representative images of Annexin V and Propidium iodide (PI) staining of uninfected and infected macrophages at 8h post infection with the indicated strains. Scale bar= 100 μM. Quantification of Annexin V (b) and PI (c) positivity at 8h post infection. Data from 2-3 experiments is represented wherein color of each circle is matched for groups within one experiment. (d and f) Percentage cell death calculated in macrophages infected at increasing MOI with H37Rv H37Rv ΔRD1, and *M. bovis* BCG after 24 hours of infection. (e and g) BODIP558/568C12 or ^14^C-oleic acid assimilation into TGs estimated by thin layer chromatography and normalized to number of cells. Data are mean ± SEM from 3 independent experiments). (h) % necrosis estimated by LDH release upon infection with H37Rv at MOI50 in the presence of indicated concentration of IM54. Data are mean ± SEM from 3 independent experiments. (i) ^14^C oleic acid assimilation into TGs estimated by thin layer chromatography and normalized to number of cells. Data are mean ± SEM from 3 independent experiments. (*p<0.05, **p<0.01, ***p<0.001). see also Figure S5.

### *Ex vivo* model of necrosis associated foamy macrophages (NAFM)

To further understand if necrosis could stimulate TG accumulation in bystander macrophages we stimulated healthy THP1 macrophages with mechanically necrotized macrophages (necrotic cell supplement/ NcS). This treatment led to accumulation of ADRP positive lipid droplets in a dose dependent manner (Figure S6). We next evaluated assimilation of ^14^C-oleate into macrophage TG, in response to NcS at a recipient to donor cell ratio of 1:2. This ratio simulated the scenario of Mtb infection at MOI10, wherein ∼67% cell death led to a lipogenic response in the remaining 33% bystander macrophages. We found a time dependent increase in exogenous FA assimilation, which was further increased in the presence of NcS (Figure 4a). Macrophages stimulated with NcS exhibited increased staining by BODIPY493/503 over a period of eight days (Figure 4b). Primary human MDMs also exhibited larger ADRP^+^ bound BODIPY493/503 stained lipid droplets (Figure 4c). Macrophages differentiated in the above manner with NcS for eight days were named necrosis associated foamy macrophages (NAFM) and normal media treated ones as normal macrophages (NM).

**Figure 4:**
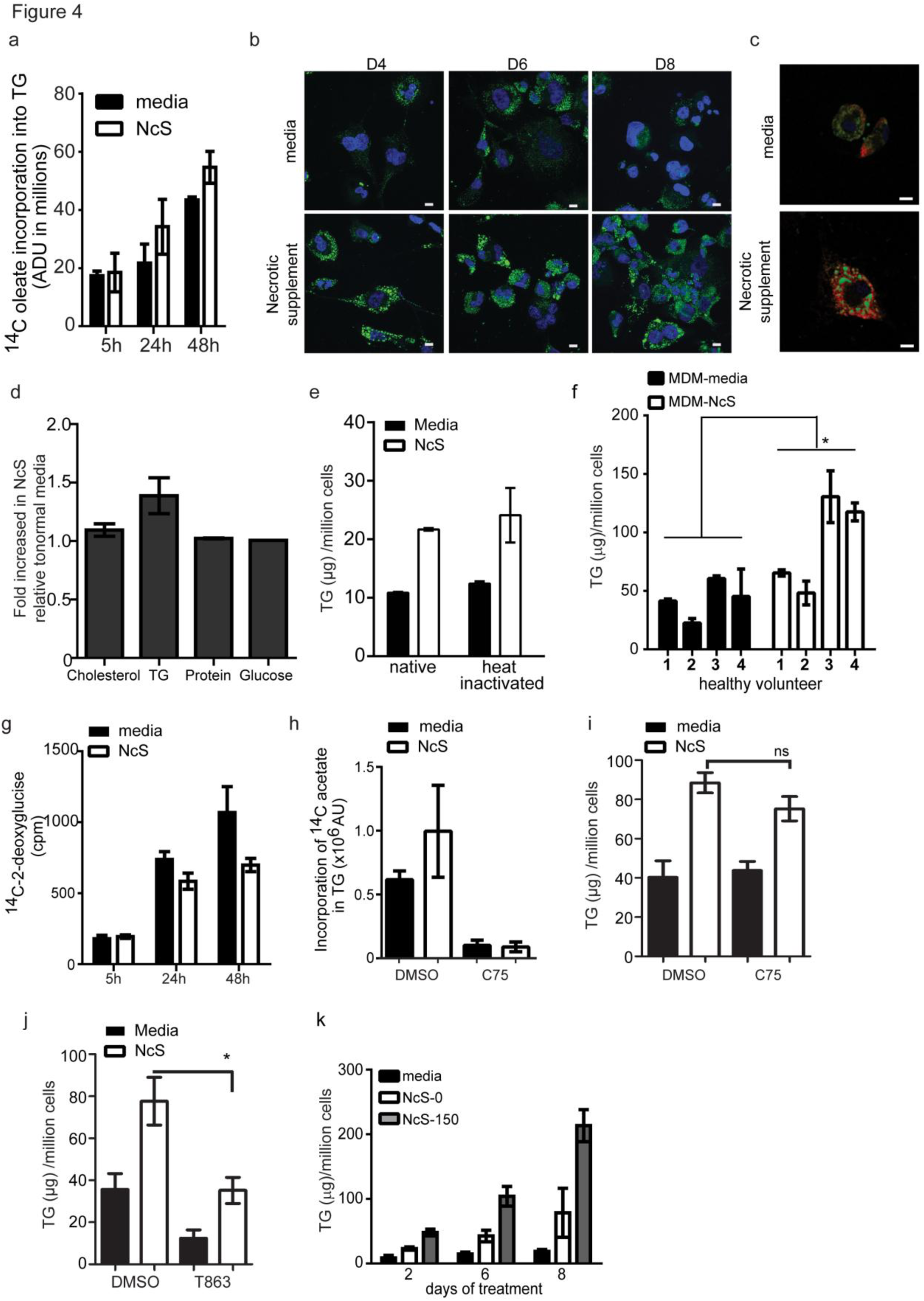
Necrotic stimulation of macrophages leads to foamy macrophage formation. (a) Incorporation of ^14^C-oleate into TG in THP1 macrophages in the presence or absence of NcS estimated by thin layer chromatography and normalized to number of cells. Data are mean ± SD from 3 wells, data representative of two independent experiments). (b) THP1 macrophages treated with media or NcS post differentiation with PMA according to the indicated time course. BODIPY493/503 (green) and DAPI (blue) stained macrophages at days 4, 6 and 8 post treatment. Scale bar=10 μM. (c) BODIPY493/503 (green) and α-ADRP (red) stained human monocyte derived macrophages (MDM) 8 days post treatment with normal media or NcS from THP1 macrophages. Scale bar=10 μM. (d) Biochemical estimation of cholesterol, TG, protein and glucose in NcS, normalized to normal growth media. Data are mean ± SD from three independent NcS preparations. (e) TG estimation eight days post differentiation with normal media, NcS or heat inactivated normal media or heat inactivated NcS. Data are mean ± SD from triplicate wells. Data are representative of 2 independent experiments. (f) TG estimation from human MDMs from 4 donors with normal media or NcS treatment over eight days. Data are mean ± SD from 3 wells, representative of three different methods of primary macrophage differentiation. Statistics between mean of media and NcS supplementation across 4 individuals are shown. (g) ^14^C-2-deocyglucose levels in macrophage cell lysates measured by scintillation counting post addition in the presence of normal media or NcS. Data are mean ± SD from triplicate wells, representative of two independent experiments. (h) Effect of C75 on *de novo* FA synthesis dependent TG levels in normal media and NcS stimulated cells. Total TG in macrophages with media or NcS after 8 days of treatment in the presence of DMSO (vehicle control), C75 (5μg/ml) (j) or T863 (10μM) (j). Data in (i) and (j) are mean ± SEM from 3 independent experiments. (ns= not statistically significant, *p<0.05). See also Figure S6. (k) TG estimation from cells post 2, 6 and 8 days of differentiation. NcS=Necrotic cell supplement. 0 and 150 indicate concentration (μM) of oleic acid used to treat THP1 macrophages prior to necrosis in normal media. Data are mean ± SD from 3 wells.

We next sought to characterize the NcS biochemically to understand the likely signals and carbon sources for this differentiation process. Cholesterol and TG in NcS increased approximately 1.1 and 1.4-fold as compared to media, while protein and glucose concentration of both media and NcS was similar (Figure 4d), suggesting a role for donor lipids in stimulating the increase in neutral lipid content of recipient cells. NcS stimulation for eight days led to 2 fold increase in TG content of recipient cells (Figure 4e). Primary human MDMs from four different donors also exhibited 1.2 to 2 fold increase in cellular TG in response to NcS over a period of eight days (Figure 4f). The involvement of a protein based signal was ruled out by heat inactivation of the NcS which failed to abrogate TG accumulation in recipient cells (Figure 4e). While exogenous FA assimilation into TG was increased, exogenous glucose uptake was in fact decreased in response to NcS (Figure 4g). This further suggested dispensability of *de novo* FA synthesis in NAFM differentiation. Inhibition of FA synthase with C75 led to 80% inhibition of *de novo* FA synthesis (Figure 4h) but did not lead to any change in total TG levels upon stimulation with NcS over an eight-day differentiation period (Figure 4i). In contrast, NcS dependent TG synthesis was abrogated by T863, an inhibitor of diacylglycerol O-acyltransferase 1 (DGAT1)^41^, verifying the role of endogenous esterification of incoming free and lipid derived FA rather than *en masse* storage of the excess exogenous TG (Figure 4j).

Unsaturated FAs such as oleic acid promote TG storage in mammalian cells ^42^; macrophages stimulated with 200 μM oleic acid increased their triglyceride content (data not shown). NcS stimulated cells increased their TG levels in a time dependent manner with a 3-5 fold increase by day 8 compared to day 0 while normal media treated cells exhibited only 1.2 fold increase in TG content during the same time (Figure 4k). NcS prepared from oleic acid stimulated macrophages further increased triglyceride levels in recipient cells by two folds in a time dependent manner (Figure 4k), suggesting a role for exogenous unsaturated fatty acids in stimulating triglyceride storage in recipient macrophages.

### TG storage in human macrophages regulates infection induced pro-inflammatory response

We questioned whether infection of NAFMs that have been differentiated by chronic exposure to necrosis, exhibit differences in immune response to Mtb infection. We found that Mtb infected NAFMs released 1.5 to 3 fold higher TNFα compared to infected NMs (Figure 5a) while neither did NAFMs exhibit a heightened basal secretion of TNFα compared to NMs (Figure 5a), nor did the NcS media itself contain detectable levels of TNFα (data not shown). We questioned if this increase was due to increased bacterial uptake in foamy macrophages. We found no difference between NM and NAFM in the uptake of Mtb (Figure 5b). Consistent with this data, we observed differences in TNFα immunostaining in necrotic granulomas compared to solid granulomas from infected guinea pig lungs (Figure 5c). TNFα seemed to be more diffuse and in general lower in case of solid granuloma sections compared to necrotic granulomas wherein TNFα expression was highest at the cuff of the necrotic center (Figure 5c).

**Figure 5:**
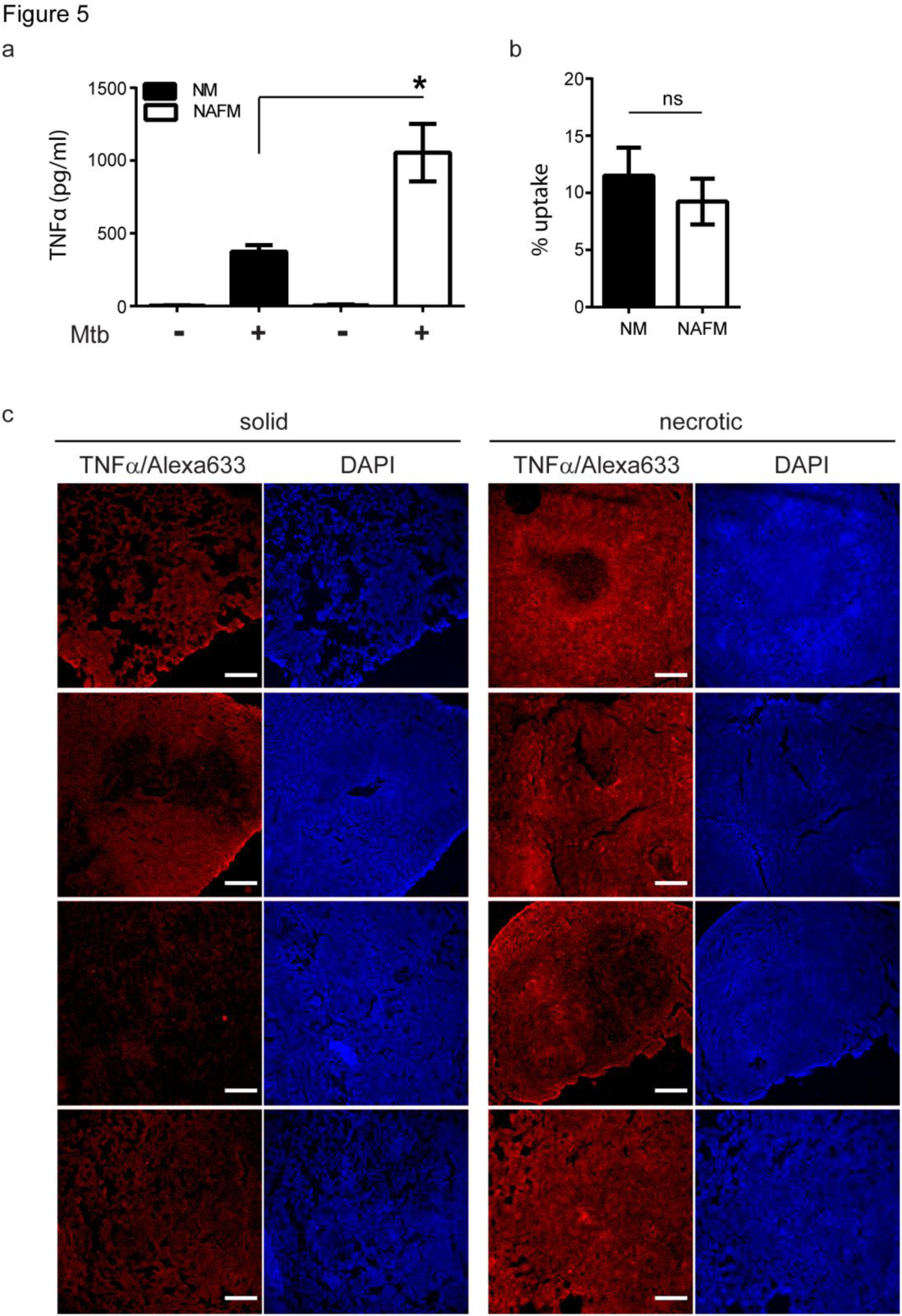
Necrosis associated foamy macrophages have a heightened inflammatory response during Mtb infection. (a) TNFα release measured by ELISA in supernatants from THP1 derived normal (NM) and necrosis associated foamy macrophages (NAFM) at 24h after infection with Mtb at MOI50 for 3h. Data are mean ± SEM from 3 independent experiments (*p≤0.05). (b) % uptake of *M. tuberculosis* in NM and NAFM after infection at MOI50 for 3h. (c) TNFα immunofluorescence in solid and necrotic guinea pig granulomas. Right panel for each section shows nuclear staining with DAPI (scale bar=250 μM).

To further test if increased cytokine production in response to infection could be regulated by TG levels, we generated two stable THP1 cell lines expressing unique shRNA against *DGAT1*. With an 80-90% knockdown in *DGAT1* expression (Figure 6a), we achieved 50-60% decrease in TG levels in THP1 macrophages (Figure 6b). Over an eight-day differentiation period, *DGAT1* knockdown cells exhibited approximately 50% decrease in TG levels in case of NMs and 30-40% decrease in case of NAFMs (Figure 6c). *DGAT1* knockdown led to approximately 50% decrease in TNFα in the supernatant (Figure 6d). To check a broader range of cytokines, chemokines, and growth factors that are expressed by macrophages in response to Mtb infection, we performed a luminex multiplex assay. We observed that besides TNFα, IL-1β, IL-1α, IL-6, GMCSF, and GCSF release upon Mtb infection was 2 to 2.5 fold higher from NAFMs compared to NMs (Figure 6e-i). In contrast, release of MCP3 was not different between infected NAFMs and NMs (Figure 6j). Infection induced release of IL-1β, IL-1α, IL-6, GMCSF, and GCSF was decreased upon depletion of *DGAT1* transcripts (Figure 6e-i). To further verify whether *DGAT1* regulated levels of these key cytokines and growth factors at the level of secretion or transcription, we quantified transcript abundance for *TNFα* and *IL1β*. Infection induced *TNFα* transcript abundance was reduced by approximately 60-80% upon *DGAT1* knockdown (Figure 6k). *IL1β* transcript abundance was also found to be 40-80% lower in NMs and 60-80% lower in NAFMs upon depletion of *DGAT1* expression (Figure 6l). Surprisingly, we found no significant differences in the uptake or growth of Mtb in NM and NAFM, whether in the wild type or *DGAT1* knockdown background (Figure S7). Therefore, intracellular TG levels regulate specific innate immune responses during Mtb infection without affecting growth of the intracellular bacilli.

**Figure 6:**
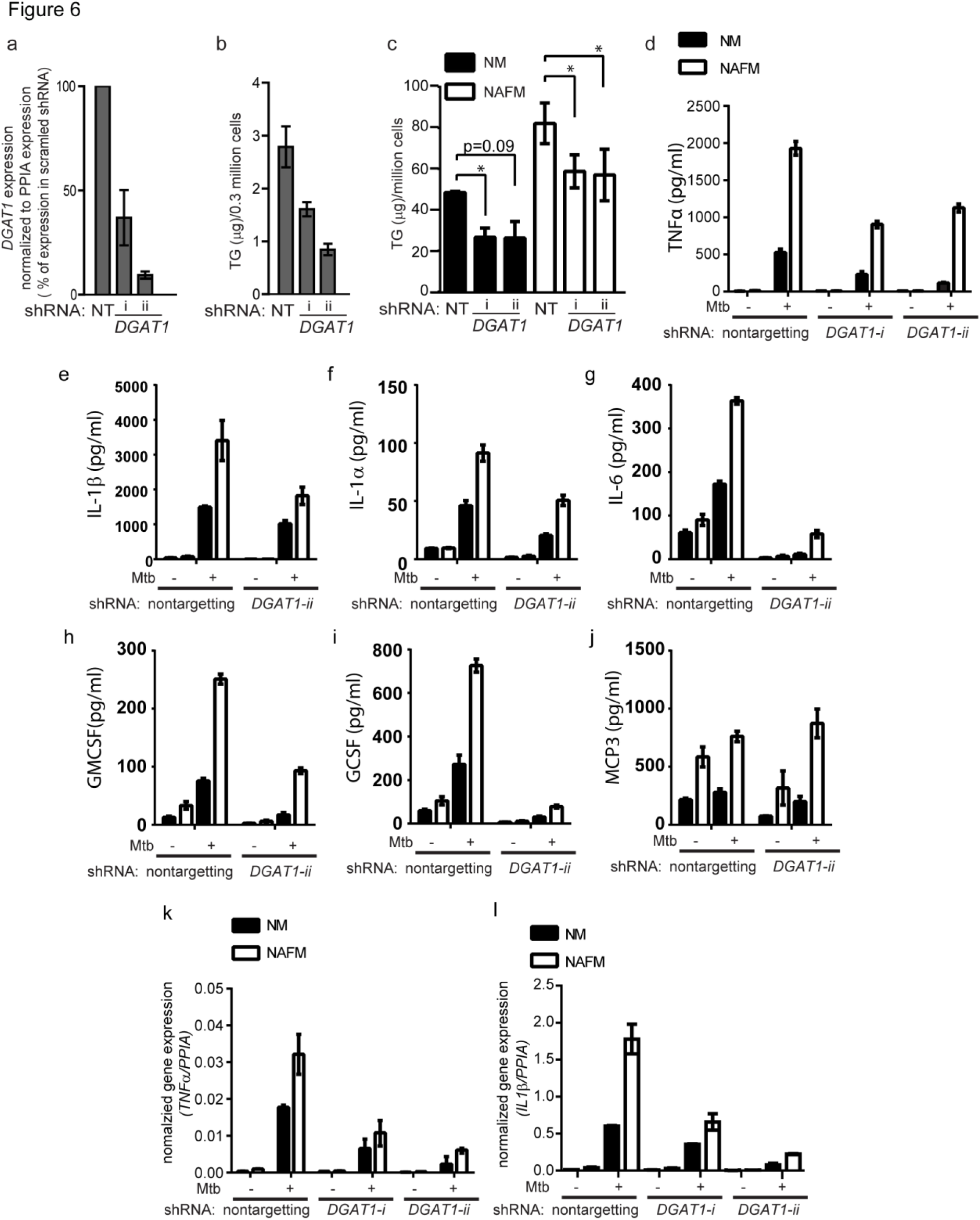
DGAT1 mediated TG synthesis is responsible for the heightened inflammatory response to Mtb infection in foamy macrophages. *DGAT1* expression (a) and TG estimation (b) in stable THP1 cell lines derived upon transduction with lentiviruses expressing non-targeting (NT) and two independent *DGAT1* targeting shRNA (i and ii). Data are mean±SD from triplicate wells (c) TG estimation at 8 days post treatment with normal media and NcS in NT and *DGAT1* knockdown THP1 macrophages after 8 days of differentiation. Data are mean ± SEM from triplicate experiments. (*p<0.05). (d) TNFα release measured by ELISA in supernatants from normal and foamy macrophages of NT and *DGAT1* knockdown THP1 macrophages at 24h after infection with Mtb at MOI50 for 3h.(Data are mean±SD from triplicate wells, representative of triplicate experiments for *DGAT1-ii* and duplicate experiments for *DGAT1-i)*. (e-j) Luminex based detection of indicated cytokines in *DGAT1-ii and Non targeting* shRNA expressing cells. Data are mean ± SEM from triplicate wells, data representative of two independent experiments. Transcript abundance of *TNFα* (k) and *IL1β* (l) normalized to that of *PPIA*. Data are mean ± SEM from triplicate wells, representative of triplicate experiments for *DGAT1-ii* and duplicate experiments for *DGAT1-i)*. See also Figure S7.

Because Mtb infection induced pro-inflammatory cytokine release in a DGAT1 dependent manner, we hypothesized that infection may be directly affecting TG metabolism. Our previous results showed that acute infection under conditions of negligible necrosis did not increase TG synthesis. To verify if TG synthesis was increased under experimental conditions used for gene expression and cytokine release measurement, we re-tested if acute infection at MOI50^3h^ followed by removal of extracellular bacteria led to alterations in TG metabolism. Synthesis of TG from exogenous and *de novo* synthesized fatty acids was measured by pulsing macrophages after infection at MOI50^3h^, with ^14^C oleic acid and ^14^C sodium acetate, in the presence of orlistat to inhibit basal lipolysis. Radioactivity incorporation into TGs was indistinguishable in infected and uninfected cells (Figure 7a and b). We next compared fatty acid assimilation over shorter durations using confocal microscopy (Figure 7c). We found that while incorporation of BODIPY558/568-C12 fatty acid into lipid droplets was comparable in infected and uninfected cells within 30 minutes of labelling, incorporation was poorer in infected cells compared to uninfected cells within 5 minutes and 15 minutes of labelling. This tracer fatty acid seemed to be accumulating more readily in bacteria compared to the host lipid droplets. This suggested that *Mtb* infection may promote lipolysis of host triglycerides rather than synthesis during acute infection wherein necrosis is not increased beyond 30% (Figure S8). To measure lipolysis, we labelled the TG pool of macrophages with ^14^C oleic acid, followed by infection at MOI50^3h^, followed by quantification of free fatty acids released in media. The addition of Triacsin C and fatty acid free BSA in the media allowed capturing even low levels of fatty acids released upon lipolysis which are otherwise undetectable. Infection led to a 2-fold and 5-fold increase in FA released at 5h and 24h post infection, respectively (Figure 7d). Therefore, infection with Mtb increased metabolism of macrophage TGs to generate free FA which may be differentially metabolised during infection.

**Figure 7:**
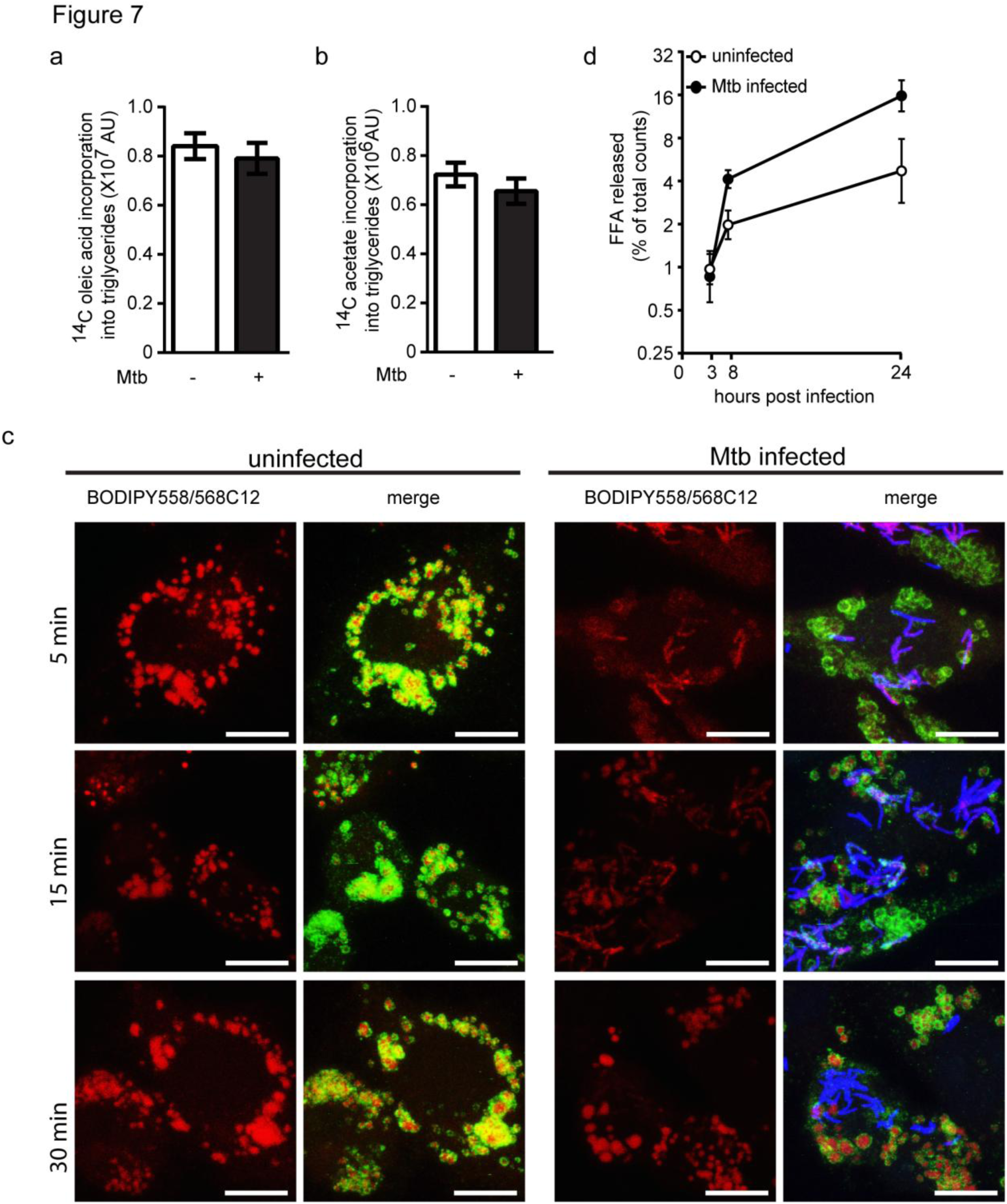
Alteration in TG metabolism upon *M. tuberculosis* infection. (a) and (b) Incorporation of ^14^C-oleate and ^14^C-acetate into TG, in the presence of orlistat, in THP1 macrophages at 21h post stopping Mtb infection at MOI50 for 3h. (c) BODIPY558/568C12 incorporation into macrophages at 21h post stopping Mtb infection at MOI50 for 3h, assessed by pulse labelling for 5, 15, and 30 min prior to fixing the macrophages. The left panel is the pseudocolored image for the FA, the right panel in each group is a merge of bacteria (blue), ADRP (green), and BODIPY558/568C12 (red). Data are representative of several frames from two independent experiments. (d) ^14^C-Free fatty acid (FFA) release measured in media of uninfected and infected cells at indicated time points post infection at MOI 50 for 3h. TG pools were ^14^C oleate labelled prior to infection. FFA release is expressed as densitometric counts of FFA fraction in media as a percentage of total densitometric counts in cellular and media fraction. Data are mean ± SD from triplicate wells representative of two experiments for (a), (b) and (c). See also Figure S8.

## Discussion

In this study, we provide evidence that TG accumulation in foamy macrophages during Mtb infection is a bystander response to necrosis both *in vitro* and *in vivo*, and that TG accumulation in macrophages renders them pro-inflammatory to Mtb infection. We provide evidence for a role of *DGAT1* in this process. These findings have important consequences on our understanding of disease etiology and macrophage differentiation. Through these investigations, we identified a novel axis of host metabolism that regulates the innate immune response to the tubercle bacillus.

Previous studies have suggested a role for oxygenated mycolic acids of mycobacteria in lipid accumulation by macrophages and monocytes during infection ^9,10^. Human macrophages cultured *in vitro* tend to store triglycerides; therefore evaluating modulation of the triglyceride content upon infection requires the use of pulse chase based assays while steady state level metabolite analysis without flux analysis may prove to be misleading. To measure synthesis, lipolysis must be inhibited and to measure lipolysis, synthesis must be inhibited ^43^. Here we used this approach to arrive at the conclusion that when human macrophages are infected with Mtb such that maximum number of cells are infected yet minimal necrosis is observed, macrophage TG indeed undergoes lipolysis while new synthesis is unaffected. We have measured flux from exogenous and *de novo* synthesized fatty acids towards TG and from TG to free fatty acids to arrive at these conclusions. In addition, we show that LipidTox series of dyes which are known to stain lipid droplets, also efficiently stain the surface of mycobacteria both *in vitro* and *in vivo* while BODIPY493/503 exclusively stains the host lipid droplets; this may be one of the reasons why our conclusions differ from a previous study that used LipidTox ^11^. Moreover, we performed an unbiased and high content image analysis on 3000-10,000 lipid droplets per experiment. Additionally, we found that the total TG abundance does not increase under infection conditions where necrosis is limiting. Dose dependent increase in MOI and the *ex vivo* model of necrosis associated foamy macrophages demonstrated that ∼67% cell death is the limit of necrosis beyond which an increase in TG content takes place in bystander macrophages. These data are consistent with the absence of oil red O positive lipid rich macrophages in solid granulomas in human tuberculosis ^10^ and in Mtb infected guinea pigs as reported by us here.

Within the heterogeneity exhibited by human TB granulomas, caseating granulomas represent a failed host immune response which is unable to restrict mycobacterial growth in a lipid rich environment ^15,16^. One of the genomic encoded regulators of necrosis is the RD1 locus of Mtb; strains deleted in this region fail to cause necrosis even in susceptible mouse models ^26,27^. RD1 region comprises of eight genes of the Esx1 secretion system, a type VII secretion system that secretes cytotoxic proteins ^27,37^. We provide evidence that upon infection with an RD1 deletion mutant of Mtb or *M. bovis* BCG, which naturally lacks this region, fatty acid assimilation as a bystander response to infection induced necrosis is prevented. It is important to note that our infection method avoids the confounding role of possible differences in growth of the three strains in the time period that we chose for our study and only reports on the effect of number of bacteria added to macrophages. Previous studies have suggested that the RD1 region in pathogenic mycobacteria is important for recruitment and activation of cells of the innate immune system *in vivo* ^44^. Our data suggest that once the cellular triglyceride levels increase, the pro-inflammatory response to Mtb also increases. The dependence on triglycerides for expression of GM-CSF and G-CSF may partly explain these defects during infection with the RD1 mutant mycobacteria.

Furthermore, pharmacological inhibition of necrosis was also able to inhibit exogenous fatty acid assimilation to TG. The inhibitor we used for this purpose, IM-54, is a mono-indolylmaleimide which is known to inhibit H_2_O_2_ mediated necrosis ^40^. The commonality between H_2_O_2_ and Mtb induced necrosis is the involvement of mitochondrial reactive oxygen species ^45^. A parallel observation has been reported in neurodegeneration using drosophila as a model; mitochondrial dysfunction and elevated ROS production in neurons has been shown to induce lipid droplet accumulation in bystander glial cells ^46^. While ROS mediated activation of SREBP was shown to be involved in this case, our data suggests increased availability of fatty acids and stimulation to increase fatty acid assimilation into triglycerides as the driver for necrosis induced triglyceride accumulation. The role of ROS in activating triglyceride accumulation may therefore be common to several diseases where cell death forms an underlying facet of the pathology.

As in case of necrosis driven foamy macrophage formation due to Mtb infection, in our *ex vivo* model of necrosis associated foamy macrophages, we found that exogenous fatty acids rather than *de novo* synthesized fatty acids provided the requisite carbon source for new TG synthesis. Unsaturated FAs stimulate TG storage ^42^; cellular lipids containing unsaturated fatty acids are liberated from necrotic cells thereby triggering the assimilation of exogenous FA into TG. This may be a generalized protective mechanism for macrophages against lipotoxicity ^42^. However, infection of these foamy macrophages with Mtb would not only ensure a safe haven for mycobacterial persistence ^47^ but as demonstrated by our data, would amplify the inflammatory response.

Lipid metabolism in macrophages plays a critical role in inflammation during metabolic disorders ^48^, atherosclerosis ^49^ and neurodegeneration ^50^. Increased TG accumulation in metabolic disorders has been reported to be majorly diet induced, where excess dietary fatty acids are stored into TG in the form of cytosolic lipid droplets ^35^. Increased inflammation in a murine model of diet induced obesity can be reversed by overexpression of DGAT1 in macrophages ^51^. Similarly increasing fatty acid degradation by overexpressing CPT1, the rate limiting enzyme in fatty acid oxidation, also plays a protective role against lipid induced inflammation ^52^. In contrast, *de novo* fatty acid biosynthesis via macrophage fatty acid synthase (FASN) has been linked with promoting the inflammatory response ^53^ and the efflux of cholesterol through ABCA1 has been linked with suppression of the inflammatory response ^54^. Our study reveals that the pro-inflammatory response of human macrophages to Mtb is tunable depending on the triglyceride content of the cells. The increase in triglyceride levels and increased synthesis of TNFα, IL-1β, IL-6, G-CSF and GM-CSF could be decreased by lowering the expressing of the last gene involved in triglyceride synthesis, *DGAT1*. We speculate that inter-individual variations in the response to *M. tuberculosis* could arise as a result of varying expression of *DGAT1*. Future studies aiming at understanding lipid storage and mobilization as modifiers of the innate response to Mtb infection would be instrumental in understanding whether this axis can be used as a potential intervention in TB treatment.

Our finding that TG hydrolysis is induced in Mtb infected cells is a step forward in understanding the mechanism for DGAT1 dependent control of inflammatory gene expression. Free fatty acids released upon TG hydrolysis act as ligands for TLR4, activating macrophages towards pro-inflammatory response ^55^. Our findings point towards a possible role of FA in mediating the inflammatory response to infection. In addition, polyunsaturated FA such as arachidonic acid or saturated FA such as palmitic acid are known mediators of inflammation ^56,57^. Arachidonic acid is particularly important in regulation of the inflammatory response in tuberculosis infection^58^ ^59^. TGs, on the other hand have been considered to be inert storage lipids, until recently when mast cells from adipose triglyceride lipase (ATGL) deficient individuals and mice were shown to be impaired in their ability to produce proinflammatory eicosanoids in response to LPS stimulation ^60^. Furthermore, LDs in which triglycerides are stored may in fact function as signalling or regulatory hubs within the cytosol. Further understanding of infection induced alterations in the flux of specific FA from TGs and alterations in the LD functional network is likely to provide mechanistic insights into TG dependent inflammatory response in tuberculosis.

A recent study reported that C57BL6 mice ^12^ exhibit presence of foamy macrophages in a TLR dependent manner, without indication of necrosis at the level of gross pathology. However, higher resolution microscopy data has revealed necrosis of macrophages and neutrophils in a burst size dependent manner during Mtb infection in C57BL6 mice ^33^. Our data from guinea pig granulomas and *in vitro* NAFM data suggests that necrosis stimulates TG accumulation in macrophages in TB and makes them pro-inflammatory. We speculate that this would be true in human TB granulomas based on recent studies revealing polarized localization of TNFα expression around the necrotic core^61^ consistent with the spatial organization of foamy macrophages in human granulomas ^10^. In the context of these studies, our data therefore suggests a two stage process in the attainment of inflammatory status as the granuloma matures: Primary recognition and recruitment of inflammatory cells which upon necrosis due to ensuing bacterial growth leads to secondary differentiation to foamy macrophages which are further pro-inflammatory to infection.

The classical necrotic granulomas are found in human post primary tuberculosis in lung tissue, or at extra-pulmonary sites such as the lymph nodes, bone, and CNS ^4,10,62^. In case of TB granulomas, TNFα has a two pronged role; early granuloma formation and control of bacterial dissemination is promoted by infection induced TNFα expression ^63,64^ while inhibiting excess TNFα expression during chronic infection is also beneficial towards control of disease ^59,65,66^. It is important to note that nonsteroidal anti-inflammatory drugs such as aspirin and ibuprofen, both of which inhibit TNFα levels, have been found to be beneficial in limiting pathology during chronic infection in the susceptible mouse model C3HeB/FeJ^8^. These approaches have illuminated the immunomodulatory role of lipid mediators in TB infection. Based on our findings, we speculate that it may be possible to regulate inflammation by targeting TG metabolism, thereby preventing host from respiratory damage and limit morbidity due to TB. Future pre-clinical studies should be aimed at evaluating the benefit of targeting TG biogenesis and mobilization in both pulmonary and extra-pulmonary disease in minimizing TB immunopathology.

## Methods

### Cell Culture and reagents

THP1 monocytes obtained from ATCC were cultured in RPMI 1640 (with glutamax, high glucose, HEPES, and sodium pyruvate) from Himedia with 10% FBS from Himedia. PMA (Phorbol 12-myristate 13-acetate), orlistat, T863 (2-((1,4-trans)-4-(4-(4-Amino-7,7-dimethyl-7H-pyrimido[4,5-b][1,4]oxazin-6-yl)- phenyl)cyclohexyl)acetic acid), and C75 (4-Methylene-2-octyl-5-oxotetrahydrofuran-3-carboxylic acid) were obtained from Sigma, TriacsinC was obtained from Alomone labs. IM54 (ALX-430-137-M001), PJ34 (ALX-270-289-M001), Necrox-2O (ALX-430-166-M001) and Necrox-5O ALX-430-247-M001) were obtained from Enzo life sciences. THP1 monocyte culture media was analysed for mycoplasma contamination using a luminescence based kit *(Lonza*, LT07-118). THP1 cell line authentication was performed at Lifecode technologies Private Limited using 10 genetic loci viz: TH01, D21S11, D5S818, D13S317, D7S820, D16S539, CSF1PO, AMEL, vWA and TPOX. The sample genotypes were queried against reference genotypes available in ATCC and DSMZ^®^ reference cell line STR databases to authenticate sample identity. *M. tuberculosis* H37Rv and *M. bovis* BCG Tokyo were kind gifts from Dr Vivek Rao, ΔRD1 and its wild type control were kind gifts of Dr David Sherman and Dr Krishnamohan Atmakuri. Mycobacterial strains were transformed with the plasmids expressing fluorescent reporter proteins. pCherry3 was a kind gift from Dr Tanya Parish (addgene#24659), pMV261-emGFP was a kind gift from Dr Vivek Rao, and pTEC18 was kind gift from Dr Lalita Ramakrishnan (addgene#30177).

### Guinea pig lung histology analysis

Guinea pigs were infected with *M. tuberculosis* H37Rv *via* the respiratory route in an aerosol chamber (Inhalation exposure system, Glas-col, IN, USA) at 100 cfu per animal. At week 4 and week 10 post infection, 3 guinea pigs were euthanized by using CO_2_ asphyxiation. After dissecting the animals, three lobes (right caudal, middle and cranial) of each lungs were fixed in 10% buffered formalin. Left caudal portion of the lung was used for the enumeration of the bacillary load. For this, the left caudal portion was weighed and homogenized in 5 ml saline in a polytron PT2100 homogenizer followed by plating on 7H11(OADC) media. Macroscopic granulomatous tissues were cut from the lung and kept in 20% sucrose solution overnight at 4°C followed by cryo-sectioning at section depth of 5μM using *Leica Cryostat CM 1850*.

*Hematoxylin and Eosin staining:* Sections on the slides were rehydrated using water for 5 min. The slides were then dipped in haematoxylin solution (1:10 Delafield’s Hematoxylin solution) for 5 min and then rinsed in water. The slides were then stain intensified by placing in ammonia solution (0.08%) for 1 min. Rinsed in water for 5 min. The sections were then equilibrated in 95% ethanol solution followed by dipping them in Eosin stain (1%) for 15 seconds. The sections were then dehydrated in 95% and 100% ethanol for 2 min in each solution, and finally rinsed in water for 5 min. The slides were then cleaned and mounted using 20% glycerol.

*Hematoxylin and oil red O staining:* The sections were first dipped in a prewarmed Oil Red O solution (0.18% in 60% isopropanol) for 1 hour at 60°C, then rinsed twice with water for 5 min each. The slides were then dipped in haematoxylin solution for 5 min, rinsed and mounted in 20% glycerol. Images were acquired using Leedz Microimaging 5MP camera attached to a Nikon Ti-U microscope. Analysis was performed by four individuals independently including a trained pathologist.

### Extraction of lipids from granulomatous lesions of Guinea pig lungs

Visible lesions from infected lungs and anatomically comparable regions from uninfected lungs were dissected out at indicated time points and then weighed. These tissues were then sonicated in 600μl of chloroform: methanol (2:1) at 60°C in a water bath sonicator. Lipids were extracted using the modified Bligh and Dyer method by first making the sonicated extract to 1200μl of chloroform: methanol (1:2) by adding 600μl of methanol. One fourth volume of 50mM Citric acid, half volume of water and one fourth volume of Chloroform was added and vortexed. This was then centrifuged at 10,000rpm for 10 min and then the lower phase taken and dried. The dried lipid extracts were weighed and then resuspended in Chloroform: Methanol (2:1) such that the concentration of the extract was 0.1 mg/μl and extracts corresponding to 0.5 mg were loaded on TLC. TLCs were developed in 4°C in the solvent system Hexane: Diethyl ether: Acetic acid (70:30:1) for neutral lipids.

### Measurement of fatty acid incorporation into TG during infection

THP-1 monocytes were differentiated into macrophages using 100 μM PMA at a density of 0.6 million cells/ml for 24 hours, followed by two days in media without PMA. Subsequently, cells were infected with *M. tuberculosis* H37Rv at respective MOI for 24h, in the presence of general lipase inhibitor orlistat ^43^ and 1μCi/ml C^14^ oleic acid *(ARC0297)* or 1 μM BODIPY558/568C12. Total cellular lipids were extracted at 24h post addition of pulse using the modified Bligh and Dyer method, dried and then analyzed using thin layer chromatography. TLCs were scanned using the Typhoon Scanner and densitometry analysis performed using Image Quant 5.2.

For confocal microscopy based experiments, macrophages were seeded on coverslips. Cells were infected at MOI 50 for 3h and 5 min, 15 min, and 30 min prior to the 24h time point, BODIPY588/568C12 was added at 1 μM to the cell culture media. At the 24h time point, media was removed, monolayers washed with pre-warmed PBS followed by fixing the monolayers with 4% neutral buffered formaldehyde. Cells were imaged using confocal microscopy.

### Lipid Extraction (modified Bligh and Dyer method)

For making a total lipid extract from THP1 monocyte derived macrophages, the cells were first washed with PBS twice and then lysed in 1% Triton X100. After lysis, 4 volumes of methanol: chloroform (2:1) was added and the lysate was vortexed. One volume of 50mM Citric acid, one volume of water and one volume of Chloroform was added and vortexed. This was then centrifuged at 10,000 rpm for 10 min, and then the lower phase isolated and dried. The dried lipid extract was then resuspended in Chloroform: Methanol (2:1) and loaded on TLC. TLCs were developed in 4°C in the solvent system Hexanes: Diethyl ether: Acetic acid (70:30:1) for neutral lipids. For visualization of unlabelled lipids on TLC, the TLCs were stained either using 10% Copper Sulphate (w/v) in 8% phosphoric acid (v/v) solution or phosphomolybdic acid (10% in ethanol) solution followed by charring at 150°C. Quantification of unlabelled lipid spots was done using ImageJ.

### Cell death assays

*Annexin V and propidium iodide staining:* THP1 monocytes were differentiated into macrophages using 100μM PMA at a density of 0.6 million cells/ml on glass coverslips for 24 hours, followed by two days in media without PMA. Macrophages were then infected with Mtb, MtbΔRD1 or *M. bovis* BCG at indicated MOI and for indicated time. Cell death by apoptosis or necrosis in macrophages infected with different strains was quantified by Alexa Flour^®^ 488 Annexin V/ Dead cell Apoptosis kit from Invitrogen. Monoloyers were washed with 1X Annexin binding buffer twice, and then stained with staining solution (10μl Alexa Flour 488 Annexin V and 1μl of 10mg/ml Propidium iodide (PI) in 100μl of 1x Annexin binding buffer) for 15 min at room temperature. Monolayers were again washed with buffer twice and fixed with 4% formaldehyde for 30 min at room temperature. The coverslips were washed with PBS and mounted on slides using DAPI mountant and sealed. Wide field Z-stacks were acquired using Leica TCS SP8 (40X objective) followed by counting annexin V or PI positive cells manually from 5 images for each group.

*Lactate Dehydrogenase release assay:* Culture supernatants from uninfected and infected THP1 macrophages were used for LDH activity assays using Cytotoxicity detection kit from Roche. Cell death was enumerated as a percentage of cells dying in treatment groups as compared to cell death in cells treated with Triton X100 (lysis buffer in kit).

*Cell counts:* THP1 macrophage monolayer was fixed with 4% formaldehyde for 15 min at room temperature followed by washing with PBS 3 times for 5 min each. Cells were then stained with 2% DAPI solution for 30 min at room temperature followed by washing with PBS 3 times for 5 min each. 10 images for each well were acquired in EVOS Floid cell imaging system using the UV lamp. Images from different groups were subjected to cell counts using Volocity software from Perkin Elmer.

### Isolation and culture of human PBMCs

Blood (9ml) was drawn from 5 individuals with their consent in K3EDTA containing vaccutainers. Blood was first diluted in RPMI to 30ml which was then layered on top of 10ml Histopaque (Sigma,10771) and centrifuged at 500g for 30min at 20°C with acceleration 9 and deceleration 1. After centrifugation the buffy coat was collected in a separate tube and washed with PBS once. To remove platelets from the pellet, the pellet was resuspended in RPMI and then layered on top of FBS followed by centrifugation. For removal of RBCs, the pellet was resuspended in 9ml of water for 10 seconds followed by addition of 1ml of 10X PBS. Cells were collected by centrifugation and then allowed to differentiate into macrophages in RPMI media containing FBS or human pooled serum, with or without human M-CSF for 6 days. In between, at day 4 the monolayer was washed to remove debris with PBS. Infections were done at day8 post isolation.

### Immunofluorescence

Cells were fixed using neutral buffered, methanol-free 4%formaldehyde, permeabilized using 0.5% Saponin, blocked with 3% bovine serum albumin (BSA) and incubated overnight with primary antibodies. Secondary antibodies used for detection were highly cross adsorbed antibodies conjugated to Alexa fluor dyes (Invitrogen). Primary antibodies used for immunostaining: CD44 (BD Pharmingen 550392), TIP47 (Progen GP30), ADRP (Progen 610102), and Perilipin (Progen GP29).

### Confocal Image analysis

THP-1 monoytes were differentiated into macrophages using 100 μM PMA at a density of 0.6 million cells/ml (on glass coverslips of 0.17mm thickness) for 24 hours, followed by two days in media without PMA. In all experiments, macrophages were fixed using 4% methanol free formaldehyde for 30 min. Fixed coverslips were washed with PBS and then stained using BODIPY493/503 (Invitrogen, D3922) solution at 10μM for one hour at room temperature, followed by three washes in 1X PBS. After staining, the coverslips were mounted on slides and sealed. Confocal z stacks of 0.30mm thickness were taken using Leica TCS SP8. Lipid droplet volume measurements were done based on BODIPY493/503 fluorescence using VOLOCITY (Perkin Elmer) image analysis software. Statistical significance was calculated from Non parametric Kruskal Walis test with a post-hoc Dunn’s test for comparison of groups. Mean lipid droplet volume from multiple experiments was tested for significance using Student’s t-test. Mean cellular fluorescence was calculated using the Leica LAX 3.1.1 software; cell boundaries were made using a free hand selection tool and total BODIPY493/503 fluorescence in the selected region of interest reported. Mean fluorescence/cell from each experiment was taken and data from 4 such experiments was pooled to derive mean and SEM. Statistical significance was calculated using Student’s t-test.

### THPM Necrosis and media preparation

Cells were scraped in media and then frozen at −20°C temperature followed by thawing at room temperature. This was repeated 5 times to make the necrotic cell supplement (NcS). NcS was then added on THP1 monocyte derived macrophages at a density of 1.2 million necrosed cells/ml media for 8 days, changing media every two days with PMA addition at day1 and day5 post addition of NcS. The“normal media” control group was differentiated in the same manner except necrotic cells were not added as supplement.

### Biochemical estimations

Quantification of Cholesterol, TG, and protein concentration in necrotic supplement and normal media was performed by specific colorimetric reactions for each of the analytes using COBAS INTEGRA 400 plus (Roche Diagnostics). Glucose estimation was performed using the anthrone method.

### Glucose uptake assays

14C −2-deoxyglucose (ARC 0112A) (1μCi/ml) was added to differentiated THP1 macropahges in media or NcS. Uptake was analyzed at 5, 24, and 48h post labelling using Perkin Elmer TopNXT Scintillation counter. Cell lysates at each time point were added to 100μl of scintillation cocktail *(Microscint PS, Perkin Elmer, 6013631)* in a white 96 well flat bottom plate and then read in the scintillation counter. Glucose uptake was plotted as ratio of radioactivity present in the cell lysate to the total radioactivity added.

### Immunofluorescence of Guinea pig lung sections

Serial sections to those used for oil red O staining were used for TNFα immunostaining. Sections were blocked with 5% BSA for 1h, followed by overnight incubation with anti-TNFα antibodies (1:50, ab1793). Alexa-633 tagged secondary antibody was used for detecting the primary antibody reactivity, followed by DAPI counterstaining. Images were acquired using a laser scanning microscope in the widefield mode.

### RNA Isolation and qRT PCR analysis

1.2 million cells were used for isolation of RNA from a single replicate of a condition, with three replicate wells per experiment. Cells were infected at MOI50 for 3h. The monolayers were washed 3 times with media and replaced with RPMI media supplemented with 10% FBS. At 24h post infection, cells were scraped in TRIzol (Ambion) or RNAzolRT (Sigma). RNA was extracted into the aqueous phase, precipitated and further purified using the Qiagen RNeasy kit or by organic precipitation. The purified RNA was DNase treated using Turbo DNAse *(Ambion, AM2238)for* 1h at 37°C. 1 μg of purified RNA was used for cDNA synthesis *(Invitrogen, 18080-093)*. cDNA was diluted 10 times and 2μl of the diluted cDNA was used for expression analysis of selected genes by qRT-PCR *(Roche, LightCycler Sybr green master mix, 04707516001)*. Primer sequences used for qRT-PCR were as follows:

PPIA_qRTF ATGCTGGACCCAACACAAAT

PPIA_qRTR TCTTTCACTTTGCCAAACACC

DGAT1_qRTF GCCTGCAGGATTCTTTCTTC

DGAT1_qRTR AGACATTGGCCGCAATAACC

TNF-α_qRTF CCCCAGGGACCTCTCTCTAATC

TNF-α_RTR GGTTTGCTACAACATGGGCTACA

IL-1β_RTF CTCGCCAGTGAAATGATGGCT

IL-1β_RTR GTCGGAGATTCGTAGCTGGAT

### *DGAT1* knockdown

Lentiviral particles for *DGAT1* knockdown were obtained from Transomic technologies (shRNA sequences used DGAT1 pZIP-mEF1 alpha zsgreen-i:

TGCTGTTGACAGTGAGCGCCCAGGTGGTGTCTCTGTTTCATAGTGAAGCCACAGATGTATGAAACAGAGACACCACCTGGATGCCTACTGCCTCGGA;

DGAT1 pZIP-mEF1 alpha zsgreen-ii: TGCTGTTGACAGTGAGCGCCCTACCGGGATGTCAACCTGATAGTGAAGCCACAGATGTATCAGGTTGACATCCCGGTAGGATGCCTACTGCCTCGGA, Non targeting control:

TGCTGTTGACAGTGAGCGAAGGCAGAAGTATGCAAAGCATTAGTGAAGCCACAGATGTAATGCT TTGCATACTTCTGCCTGT GCCTACTGCCTCGGA). 50,000 THP1 monocytes were transduced with lentiviral particles at MOI10 for 24h. Transduced cell lines were maintained for stable cell line generation in puromycin at 0.6μg/ml for about 3-4 weeks. Knockdown efficiency was checked using qRT-PCR and also validated using TG analysis from TLC.

### Cytokine analysis

THP1 macrophages were differentiated as described above for 8 days, followed by infection with *M. tuberculosis* at MOI50 for 3 hours at which point extracellular bacilli were removed. The monolayers were washed 3 times with media and replaced with RPMI media supplemented with 10% FBS. Cell culture supernatants were harvested at 24h post infection and assayed for TNFα using an ELISA kit (eBioscience) or Milliplex human cytokine/chemokine bead panel (HCYTOMAG-60K).

### Fatty acid release assays

C14-oleic acid was added to differentiated THP1 macrophages (normal or foamy differentiated for 8 days) to label macrophage TG for 16h. After removal of extracellular label (washing with media 4 times), macrophages were infected at MOI50 for 3h. Extracellular bacteria were washed off by washing with PBS 4 times and then fatty acid free BSA medium was added (2% fatty acid free BSA in RPMI) along with Triacsin C (10μM). Total lipids were extracted from culture supernatants and cell lysates separately at different time points post addition of Triacsin C. Lipids were resolved by thin layer chromatography as described above. Densitometry was performed using Image Quant 5.2. Fatty acid released or TG remaining was plotted as ratio of densitometric units present in fatty acid band to the total density in cell lysate and supernatant at that time point.

### Intracellular mycobacterial growth assay

Macrophages were seeded in 48 well plates at a density of0.6 million/ml and infected at MOI of 0.1 for 4 hours, followed by removal of extracellular bacilli by 3 washed with PBS and addition of complete growth media. Cells were lysed in 100ul of 1% Triton-X100 and serial dilutions plated on 7H10 OADC plates. Media was changed after every 48 hours.

### Statistics

All statistics were computed using Prism. ANOVA and t-test were used for all parametric data wherein data from 3-4 independent experiments were combined. Only for testing statistical significance of change in median of the lipid droplet size (Fig.2c), Kruskal Wallis test was used.

### Ethics Statement

The animals were housed and handled at the Department of Biochemistry, University of Delhi South Campus according to directives and guidelines of the Committee for the Purpose of Control and Supervision of Experiments on Animals (CPCSEA). Use of *M. tuberculosis* infected guinea pig lung tissue for this work was approved by the CPCSEA as per ethics proposal #IGIB/IAEC/14/15. Blood was drawn from healthy human volunteers with informed written consent as per approval #11dtd.March30th2015 of the Institutional Human Ethics Committee.

## Abbreviations

Triglyceride (TG), Fatty acid (FA), Diacylglycerol O-acyltransferase 1 (DGAT1), Necrotic cell supplement (NcS), Necrosis associated foamy macrophages (NAFM), normal macrophages (NM).

## Acknowledgements

This study is funded by Wellcome Trust-DBT India Alliance (Grant IA/I/11/2500254) to SG. SG acknowledges BSL3 facility (STS0016) and imaging facility (BSC0403) support by Council of Scientific and Industrial Research. Authors thank DBT, CSIR, and UGC for PhD fellowships to SD (DBT), KS (CSIR), AN and PDB (UGC).The authors thank Vivek Rao and Rajesh S Gokhale for constructive comments and suggestions during the course of the research work. The authors thank Mr Manish Kumar for maintenance of the imaging facility.

## Author contributions

Conceptualization: SG; Methodology: SG and NJ; Investigation: NJ and SD; Microscopy: NJ, SD, KS, AN and SG; Validation: DM, KS, AN, and PDB; Analysis: NJ and SG; Resources: GK, AKT, SG; Writing: NJ and SG; Supervision: SG; Funding Acquisition: SG

## Supplemental Information

**Figure S1.**
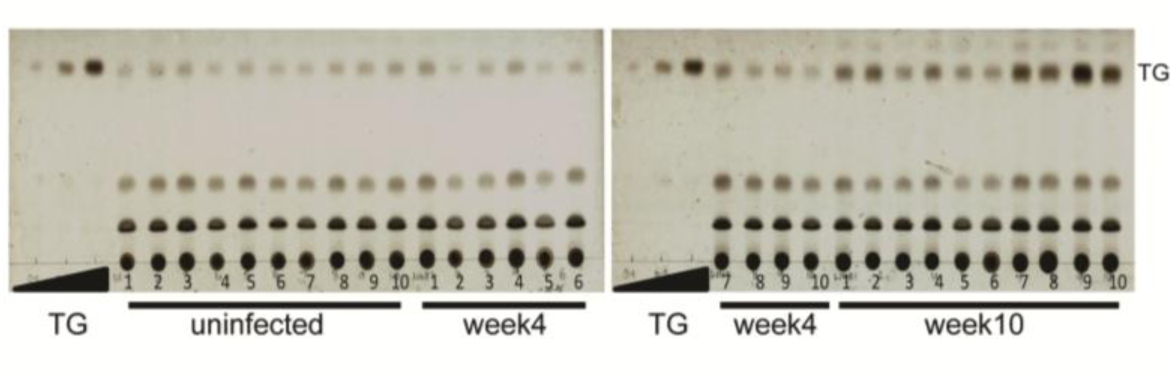
Related to Figure 1. Guinea pig granulomatous tissues containing necrotic lesions are rich in TGs. TLC for TG estimation in 10 sections each from lungs of uninfected, week4 infected and week 10 infected guinea pigs.

**Figure S2.**
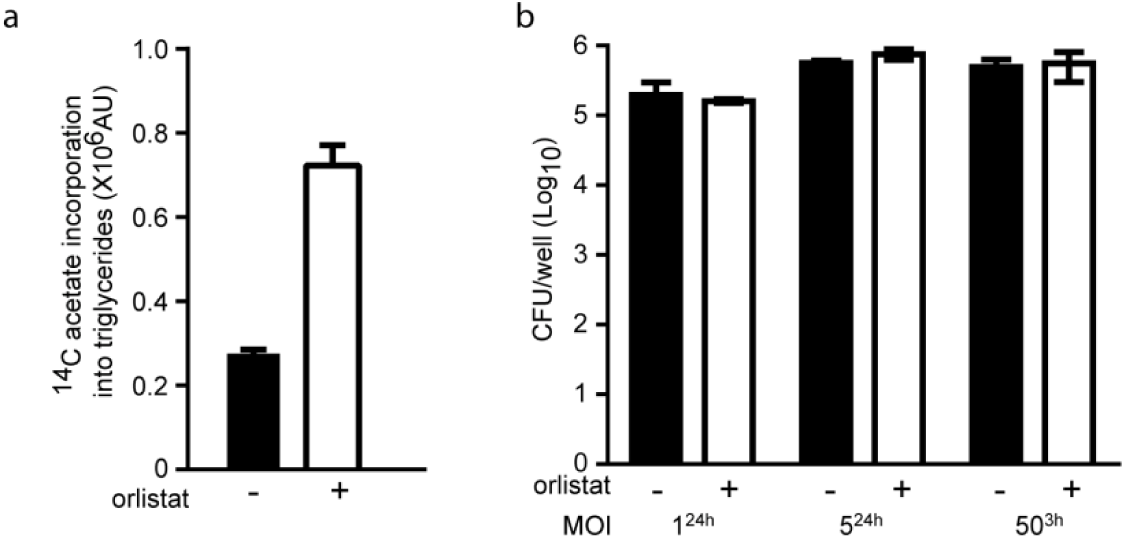
Related to Figure 2 and 3. Inhibition of fatty acid hydrolysis by Orlistat allows measurement of absolute rate of TG synthesis. (a) ^14^C-acetate incorporation in TG measured using thin layer chromatography post treatment in the presence of 10 μM Orlistat or vehicle control (DMSO) for 24 hours. (b) Bacterial viability in macrophages treated with Orlistat at 10 μM at 24h post infection at 24h 24 3h time for which bacteria were added during infection MOI1, MOI5, MOI50 (MOI). All groups were sampled at the same time i.e. 24h post addition of bacteria. In case of MOI 50^3h^ extracellular bacteria were removed and monolayers washed at 3h and then treated with DMSO or orlistat for 21 hours.

**Figure S3.**
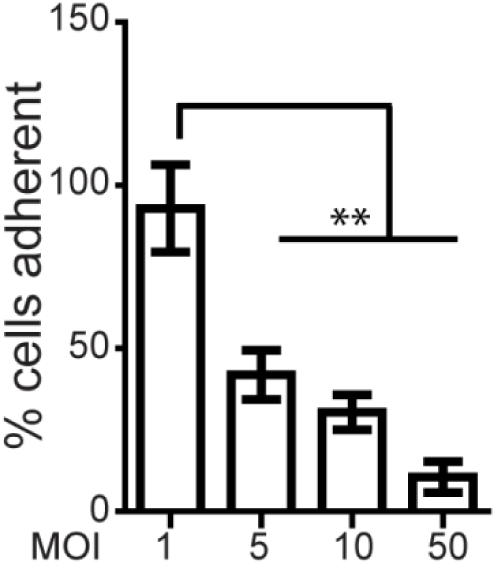
Related to Figure 2. Necrosis during infection. Graph represents percentage of cells remaining adherent at 24h post infection at indicated MOI as a percentage of uninfected control. Data are mean ± SEM from 3 independent experiments. (p values indicated and as per **p<0.01)

**Figure S4.**
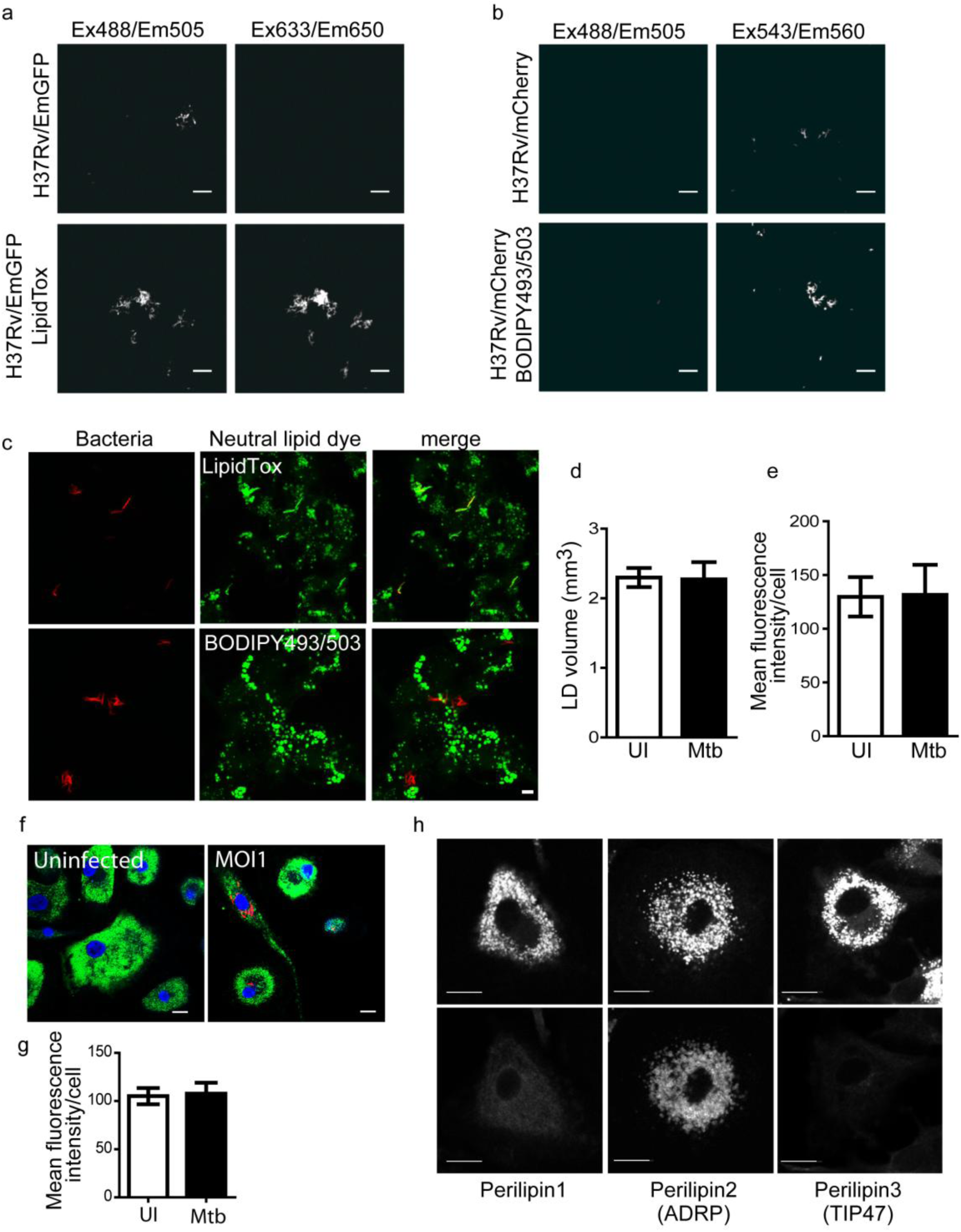
Related to Figure 2. Imaging based assessment of neutral lipid content in human monocyte derived macrophages. (a and b) LipidTox Deep Red and BODIPY 493/503 (lower panels) staining of *in vitro* cultured *M. tuberculosis* H37Rv expressing emGFP and mCherry respectively. Scale bar=5 μm. Monochrome images have been shown. (c) LipidTox Deep Red (upper middle panel) and BODIPY 493/503 (lower middle panel) staining of THP1 macrophages infected with *M. tuberculosis* H37Rv expressing emGFP (top left panel) and mCherry (bottom left panel) respectively. Signal in the channel for bacteria has been pseudocolored red and the signal for the lipid stain has been pseudocolored green for representation. Scale bar=5 μm. Mean lipid droplet volume (d) and mean fluorescence BODIPY493/503 intensity per cell (e) of uninfected and mCherry+ Mtb infected THP1 macrophages after 24h of infection at MOI1. Data are mean±SEM from 4 and 3 independent experiments respectively. (f) Perilipin1, ADRP, and TIP47 immunostaining (lower panels) and lipid droplet staining (BODIPY 493/503, upper panel) in human monocyte derived macrophages reveals ADRP specific coating of lipid droplets in human MDMs. (g) BODIPY 493/503 staining of uninfected and mCherry+*M. tuberculosis* infected human monocyte derived macrophages. Scale bar=5 μm. (h) Mean BODIPY493/503 fluorescence per cell in uninfected and mCherry^+^ *M. tuberculosis* infected human monocyte derived macrophages after 24h of infection at MOI1. Data is mean ± SD from four healthy individuals. Scale bar=10μm.

**Figure S5.**
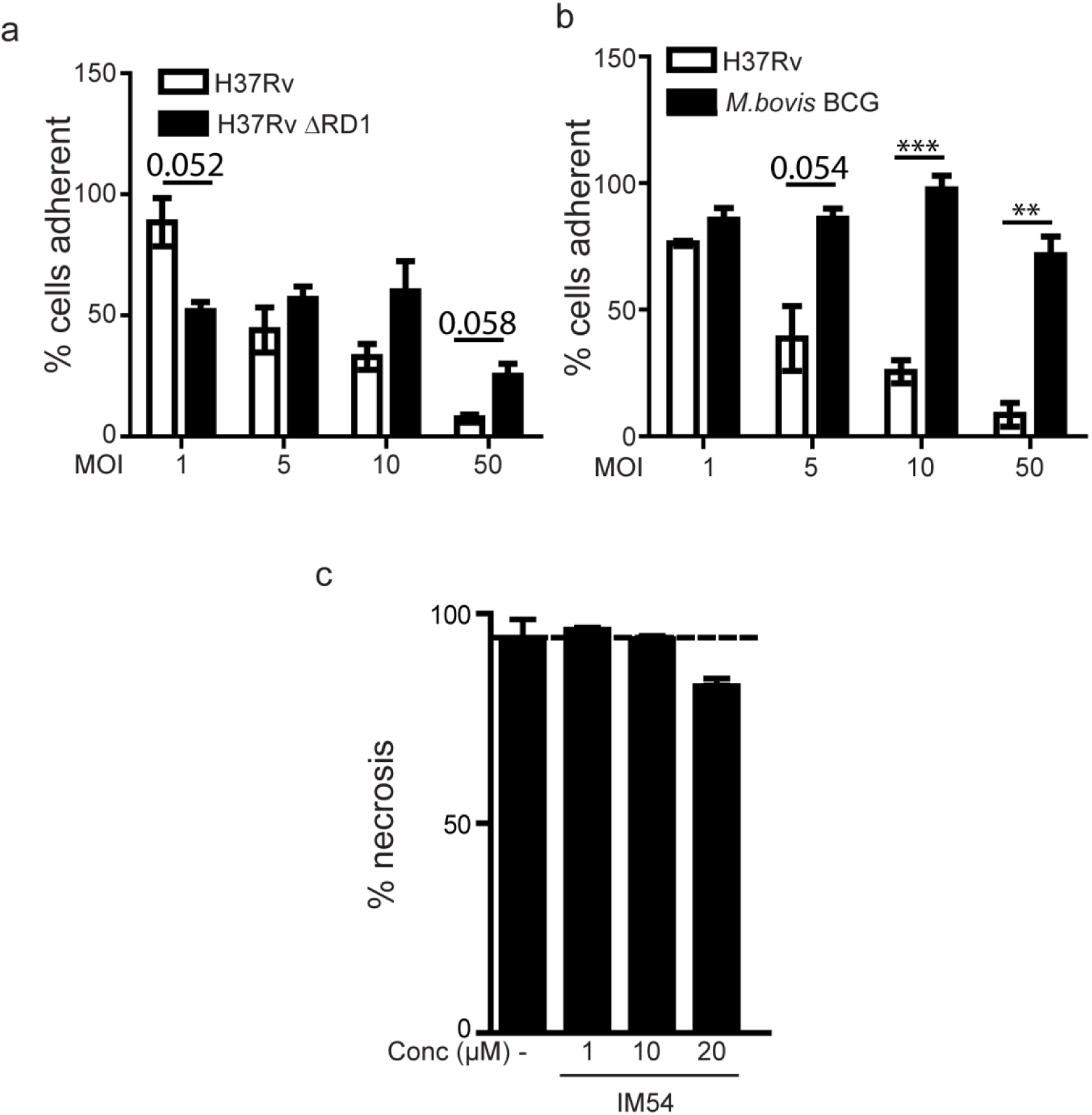
Related to Figure 3. Necrosis and TG synthesis during infection. (a and b) Graph represents percentage of cells remaining adherent at 24h post infection at indicated MOI as a percentage of uninfected control. Data are mean ± SEM from 3 independent experiments. (p values indicated and as per **p<0.01, **p<0.001) (c) Graph represents cell death as % necrosis (from cell counts) in MOI50 *M. tuberculosis* infected THP1 macrophages in the presence of increasing concentrations of indicated necrosis inhibitors. Dotted line indicates necrosis in MOI50^24h^ condition. Data are mean ± SD from triplicate wells.

**Figure S6.**
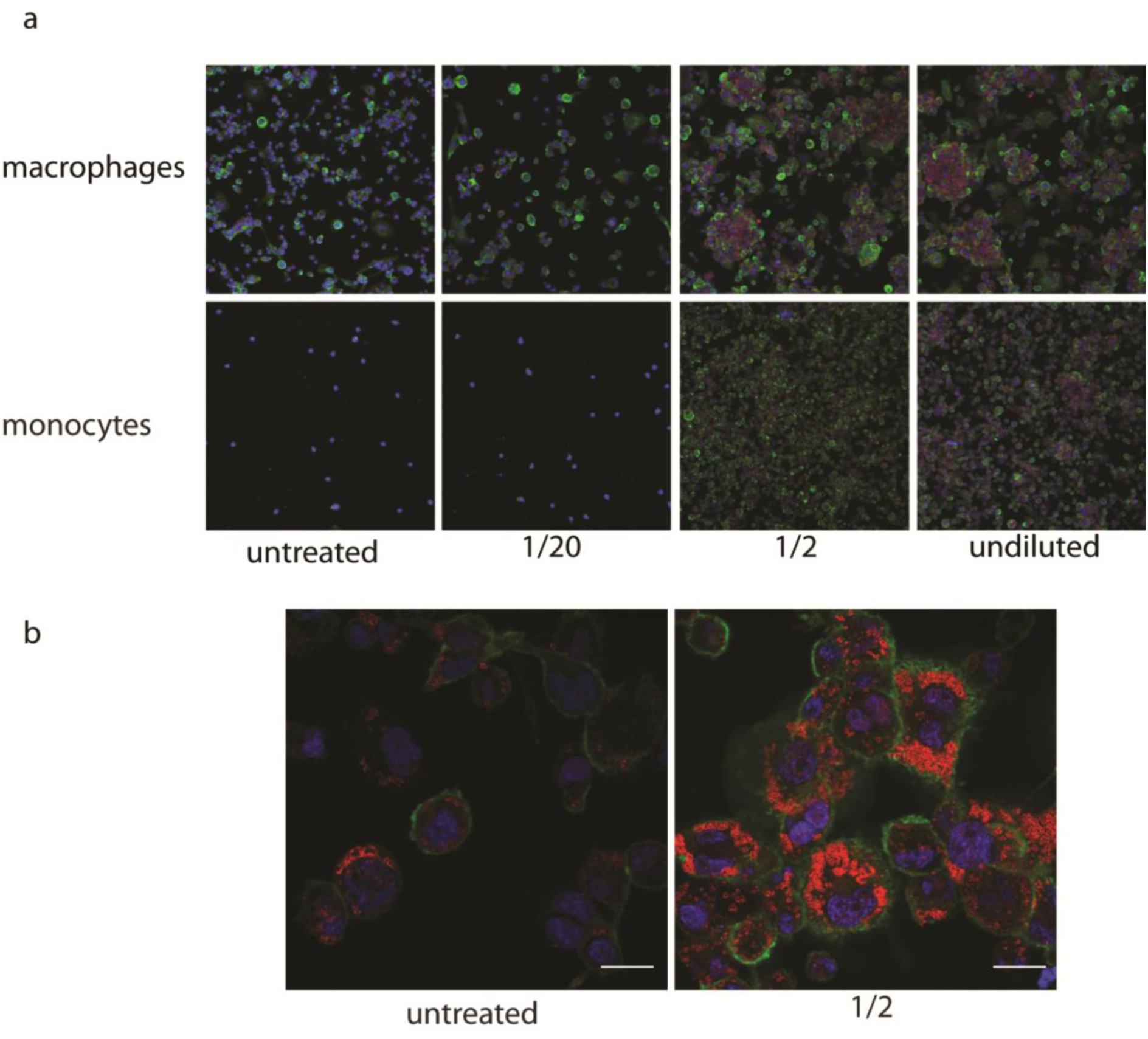
Related to Figure 4. Dosage effect of necrotic cell stimulation towards accumulation of ADRP positive lipid droplets in THP-1 macrophages and monocytes. (a). Low magnification images of macrophages (top panel) and monocytes (bottom panel) that were treated with indicated dilutions of necrotic supplement in normal media for 2 days. Cells are stained for CD44 (green) to mark cell surface of macrophages and ADRP (red) to mark the lipid droplets. Nuclei were stained using DAPI (blue). (b) High magnification imaging reveals high density of intracellular, ADRP positive organelles in macrophages stimulated with necrotic supplement. Scale bar= 10 μM.

**Figure S7.**
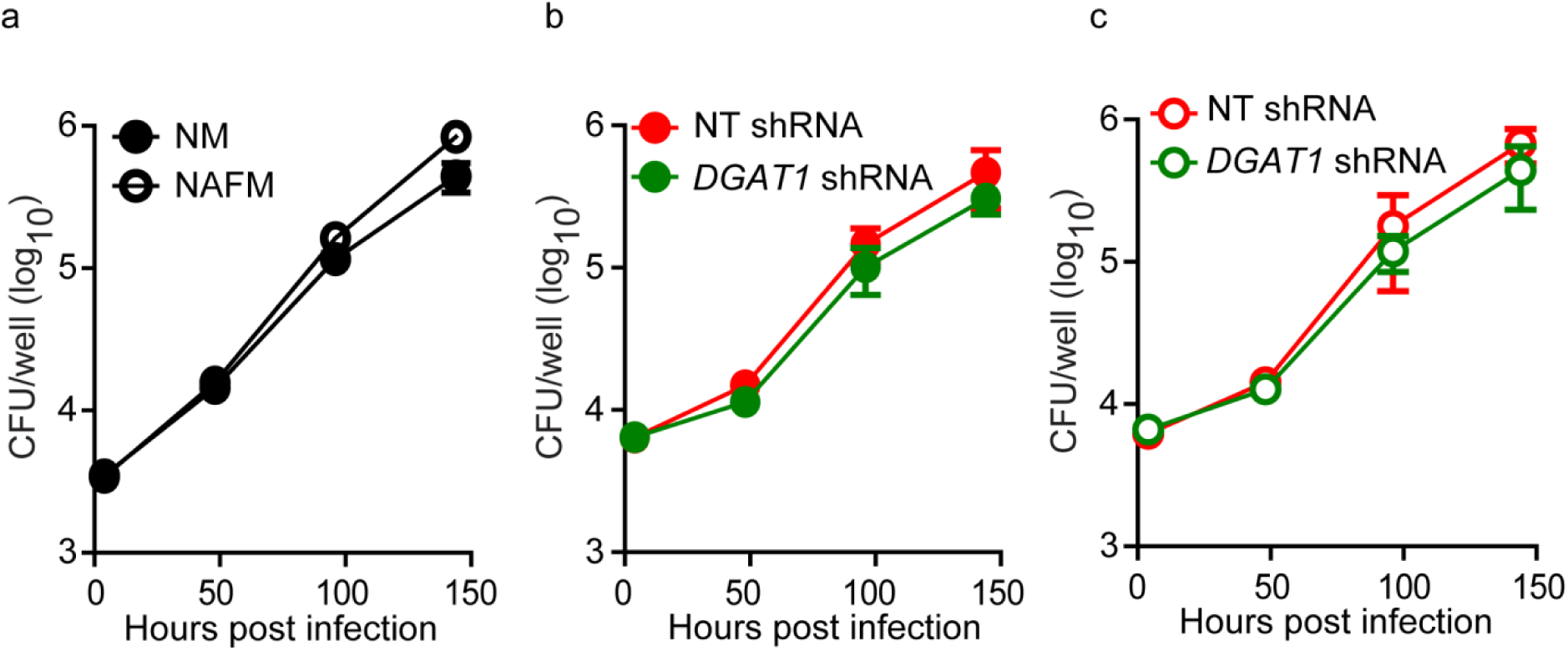
Related to Figure 6. Uptake and growth of *M. tuberculosis* in NM and NAFM post infection for 4h at MOI 0.1. (a) represents intracellular growth of *M. tuberculosis* H37Rv in NM and NAFM. (b) represents intracellular growth of *M. tuberculosis* H37Rv in NM from non-targeting shRNA and DGAT1 shRNA expressing macrophages (shRNA-ii). (c) represents intracellular growth of *M. tuberculosis* H37Rv in NAFM from non-targeting shRNA and DGAT1 shRNA expressing macrophages (shRNA-ii). Data are mean ± SD from triplicate wells. Data are representative of 3 experiments for (a) and 2 experiments for (b) and (c).

**Figure S8.**
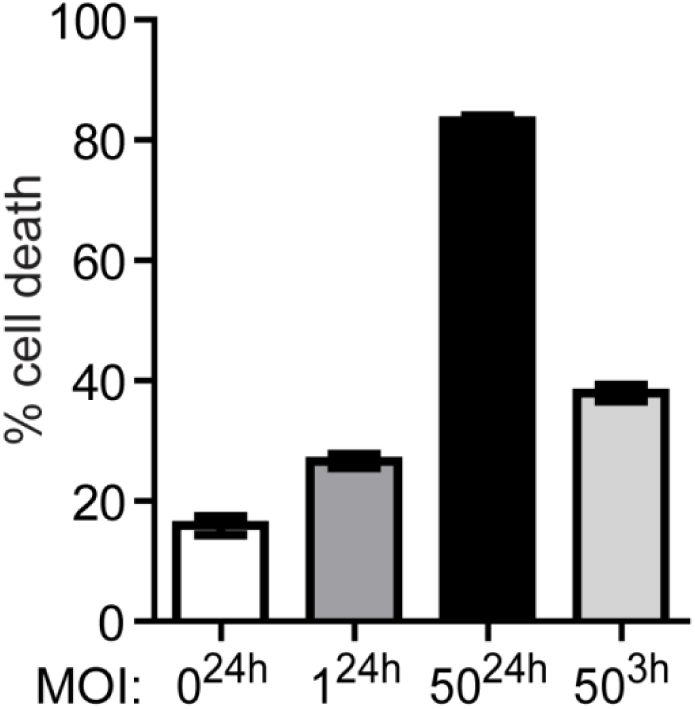
Related to Figure 7. Cell death in an acute model of infection. Cell death quantified by LDH release in uninfected and Mtb H37Rv infected THP1 macrophages, presented as % of Triton X100 lysed macrophages. 1^24h^, 50^24h^, 50^3h^ indicate the MOI^time for which bacteria were added during infection^. All groups were sampled at the same time i.e. 24h post addition of bacteria. In case of MOI 50^3h^ extracellular bacteria were removed and monolayers washed at 3h and replaced with fresh medium. Data are mean ± SD of triplicate wells.

## References

1. WHO report. www.who.int/mediacentre/factsheets/fs310/en/ (2015).

2. Bloch, H. & Segal, W. Biochemical differentiation of Mycobacterium tuberculosis grown in vivo and in vitro. J Bacteriol 72, 132–141 (1956).

3. Ehrt, S., Rhee, K. & Schnappinger, D. Mycobacterial genes essential for the pathogen’s survival in the host. Immunol Rev 264, 319–326 (2015).

4. Kim, M. J. et al. Caseation of human tuberculosis granulomas correlates with elevated host lipid metabolism. EMBO Mol Med 2, 258–274 (2010).

5. Wallis, R. S. & Hafner, R. Advancing host-directed therapy for tuberculosis. Nat Rev Immunol 15, 255–263 (2015).

6. Korf, H. et al. Liver X receptors contribute to the protective immune response against Mycobacterium tuberculosis in mice. J Clin Invest 119, 1626–1637 (2009).

7. Bouttier, M. et al. Alu repeats as transcriptional regulatory platforms in macrophage responses to M. tuberculosis infection. Nucleic Acids Res 44, 10571–10587 (2016).

8. Vilaplana, C. et al. Ibuprofen therapy resulted in significantly decreased tissue bacillary loads and increased survival in a new murine experimental model of active tuberculosis. J Infect Dis 208, 199–202 (2013).

9. Dkhar, H. K. et al. Mycobacterium tuberculosis keto-mycolic acid and macrophage nuclear receptor TR4 modulate foamy biogenesis in granulomas: a case of a heterologous and noncanonical ligand-receptor pair. J Immunol 193, 295–305 (2014).

10. Peyron, P. et al. Foamy macrophages from tuberculous patients’ granulomas constitute a nutrient-rich reservoir for M. tuberculosis persistence. PLoS Pathog 4, e1000204 (2008).

11. Singh, V. et al. Mycobacterium tuberculosis-driven targeted recalibration of macrophage lipid homeostasis promotes the foamy phenotype. Cell Host Microbe 12, 669–681 (2012).

12. Holla, S. et al. MUSASHI-Mediated Expression of JMJD3, a H3K27me3 Demethylase, Is Involved in Foamy Macrophage Generation during Mycobacterial Infection. PLoS Pathog 12, e1005814 (2016).

13. Underhill, D. M., Ozinsky, A., Smith, K. D. & Aderem, A. Toll-like receptor-2 mediates mycobacteria-induced proinflammatory signaling in macrophages. Proc Natl Acad Sci U S A 96, 14459–14463 (1999).

14. Drennan, M. B. et al. Toll-like receptor 2-deficient mice succumb to Mycobacterium tuberculosis infection. Am J Pathol 164, 49–57 (2004).

15. Cadena, A. M., Fortune, S. M. & Flynn, J. L. Heterogeneity in tuberculosis. Nat Rev Immunol July 24 (2017).

16. Orme, I. M., Robinson, R. T. & Cooper, A. M. The balance between protective and pathogenic immune responses in the TB-infected lung. Nat Immunol 16, 57–63 (2015).

17. Martin, C. J. et al. Efferocytosis is an innate antibacterial mechanism. Cell Host Microbe 12, 289–300 (2012).

18. Behar, S. M. et al. Apoptosis is an innate defense function of macrophages against Mycobacterium tuberculosis. Mucosal Immunol 4, 279–287 (2011).

19. Lee, J., Repasy, T., Papavinasasundaram, K., Sassetti, C. & Kornfeld, H. Mycobacterium tuberculosis induces an atypical cell death mode to escape from infected macrophages. PLoS One 6, e18367 (2011).

20. Duan, L., Gan, H., Golan, D. E. & Remold, H. G. Critical role of mitochondrial damage in determining outcome of macrophage infection with Mycobacterium tuberculosis. J Immunol 169, 5181–5187 (2002).

21. Divangahi, M., Desjardins, D., Nunes-Alves, C., Remold, H. G. & Behar, S. M. Eicosanoid pathways regulate adaptive immunity to Mycobacterium tuberculosis. Nat Immunol 11, 751–758 (2010).

22. Cooper, A. M. Cell-mediated immune responses in tuberculosis. Annu Rev Immunol 27, 393–422 (2009).

23. O’Garra, A. et al. The immune response in tuberculosis. Annu Rev Immunol 31, 475–527 (2013).

24. Menezes, A. M. et al. Tuberculosis and airflow obstruction: evidence from the PLATINO study in Latin America. Eur Respir J 30, 1180–1185 (2007).

25. Hnizdo, E., Singh, T. & Churchyard, G. Chronic pulmonary function impairment caused by initial and recurrent pulmonary tuberculosis following treatment. Thorax 55, 32–38 (2000).

26. Junqueira-Kipnis, A. P. et al. Mycobacteria lacking the RD1 region do not induce necrosis in the lungs of mice lacking interferon-gamma. Immunology 119, 224–231 (2006).

27. Simeone, R. et al. Phagosomal rupture by Mycobacterium tuberculosis results in toxicity and host cell death. PLoS Pathog 8, e1002507 (2012).

28. Conrad, W. H. et al. Mycobacterial ESX-1 secretion system mediates host cell lysis through bacterium contact-dependent gross membrane disruptions. Proc Natl Acad Sci U S A 114, 1371–1376 (2017).

29. Kinhikar, A. G. et al. Potential role for ESAT6 in dissemination of M. tuberculosis via human lung epithelial cells. Mol Microbiol 75, 92–106 (2010).

30. McMurray, D. N. in Tuberculosis (American Society of Microbiology, 1994).

31. Turner, O. C., Basaraba, R. J. & Orme, I. M. Immunopathogenesis of pulmonary granulomas in the guinea pig after infection with Mycobacterium tuberculosis. Infect Immun 71, 864–871 (2003).

32. Hoff, D. R. et al. Location of intra-and extracellular M. tuberculosis populations in lungs of mice and guinea pigs during disease progression and after drug treatment. PLoS One 6, e17550 (2011).

33. Repasy, T. et al. Intracellular bacillary burden reflects a burst size for Mycobacterium tuberculosis in vivo. PLoS Pathog 9, e1003190 (2013).

34. Palmer, A. M. et al. Differential uptake of subfractions of triglyceride-rich lipoproteins by THP-1 macrophages. Atherosclerosis 180, 233–244 (2005).

35. Farese, R. V., Jr. & Walther, T. C. Lipid droplets finally get a little R-E-S-P-E-C-T. Cell 139, 855–860 (2009).

36. Smith, J. et al. Evidence for pore formation in host cell membranes by ESX-1-secreted ESAT-6 and its role in Mycobacterium marinum escape from the vacuole. Infect Immun 76, 5478–5487 (2008).

37. Mahairas, G. G., Sabo, P. J., Hickey, M. J., Singh, D. C. & Stover, C. K. Molecular analysis of genetic differences between Mycobacterium bovis BCG and virulent M. bovis. J Bacteriol 178, 1274–1282 (1996).

38. Dobos, K. M., Spotts, E. A., Quinn, F. D. & King, C. H. Necrosis of lung epithelial cells during infection with Mycobacterium tuberculosis is preceded by cell permeation. Infect Immun 68, 6300–6310 (2000).

39. Riendeau, C. J. & Kornfeld, H. THP-1 cell apoptosis in response to Mycobacterial infection. Infect Immun 71, 254–259 (2003).

40. Dodo K, K. M., Shimizu T, Takahashi M, Sodeoka M. Inhibition of hydrogen peroxide-induced necrotic cell death with 3-amino-2-indolylmaleimide derivatives. Bioorganic and Medicinal Chemistry Letters 15, 3114–3118 (2005).

41. Cao, J. et al. Targeting Acyl-CoA:diacylglycerol acyltransferase 1 (DGAT1) with small molecule inhibitors for the treatment of metabolic diseases. J Biol Chem 286, 41838–41851 (2011).

42. Listenberger, L. L. et al. Triglyceride accumulation protects against fatty acid-induced lipotoxicity. Proc Natl Acad Sci U S A 100, 3077–3082 (2003).

43. Brasaemle, D. L. et al. Perilipin A increases triacylglycerol storage by decreasing the rate of triacylglycerol hydrolysis. J Biol Chem 275, 38486–38493 (2000).

44. Volkman, H. E. et al. Tuberculous granuloma formation is enhanced by a mycobacterium virulence determinant. PLoS Biol 2, e367 (2004).

45. Roca, F. J. & Ramakrishnan, L. TNF dually mediates resistance and susceptibility to mycobacteria via mitochondrial reactive oxygen species. Cell 153, 521–534 (2013).

46. Liu, L. et al. Glial lipid droplets and ROS induced by mitochondrial defects promote neurodegeneration. Cell 160, 177–190 (2015).

47. Cardona, P. J. et al. Widespread bronchogenic dissemination makes DBA/2 mice more susceptible than C57BL/6 mice to experimental aerosol infection with Mycobacterium tuberculosis. Infect Immun 71, 5845–5854 (2003).

48. Prieur, X. et al. Differential lipid partitioning between adipocytes and tissue macrophages modulates macrophage lipotoxicity and M2/M1 polarization in obese mice. Diabetes 60, 797–809 (2011).

49. Chinetti-Gbaguidi, G., Colin, S. & Staels, B. Macrophage subsets in atherosclerosis. Nat Rev Cardiol 12, 10–17 (2015).

50. Boven, L. A. et al. Myelin-laden macrophages are anti-inflammatory, consistent with foam cells in multiple sclerosis. Brain 129, 517–526 (2006).

51. Koliwad, S. K. et al. DGAT1-dependent triacylglycerol storage by macrophages protects mice from diet-induced insulin resistance and inflammation. J Clin Invest 120, 756–767 (2010).

52. Malandrino, M. I. et al. Enhanced fatty acid oxidation in adipocytes and macrophages reduces lipid-induced triglyceride accumulation and inflammation. Am J Physiol Endocrinol Metab 308, E756–769 (2015).

53. Moon, J. S. et al. UCP2-induced fatty acid synthase promotes NLRP3 inflammasome activation during sepsis. J Clin Invest 125, 665–680 (2015).

54. Ito, A. et al. LXRs link metabolism to inflammation through Abca1-dependent regulation of membrane composition and TLR signaling. 4, e08009 (2015).

55. Lee, J. Y. et al. Saturated fatty acid activates but polyunsaturated fatty acid inhibits Toll-like receptor 2 dimerized with Toll-like receptor 6 or 1. J Biol Chem 279, 16971–16979 (2004).

56. Schilling, J. D. et al. Palmitate and lipopolysaccharide trigger synergistic ceramide production in primary macrophages. J Biol Chem 288, 2923–2932 (2013).

57. Tobin, D. M. & Ramakrishnan, L. TB: the Yin and Yang of lipid mediators. Curr Opin Pharmacol 13, 641–645 (2013).

58. Mayer-Barber, K. D. & Sher, A. Cytokine and lipid mediator networks in tuberculosis. Immunol Rev 264, 264–275 (2015).

59. Tobin, D. M. et al. The lta4h locus modulates susceptibility to mycobacterial infection in zebrafish and humans. Cell 140, 717–730 (2010).

60. Dichlberger, A., Schlager, S., Maaninka, K., Schneider, W. J. & Kovanen, P. T. Adipose triglyceride lipase regulates eicosanoid production in activated human mast cells. J Lipid Res 55, 2471–2478 (2014).

61. Marakalala, M. J. et al. Inflammatory signaling in human tuberculosis granulomas is spatially organized. Nat Med 22, 531–538 (2016).

62. Hunter, R. L., Jagannath, C. & Actor, J. K. Pathology of postprimary tuberculosis in humans and mice: contradiction of long-held beliefs. Tuberculosis (Edinb) 87, 267–278 (2007).

63. Bean, A. G. et al. Structural deficiencies in granuloma formation in TNF gene-targeted mice underlie the heightened susceptibility to aerosol Mycobacterium tuberculosis infection, which is not compensated for by lymphotoxin. J Immunol 162, 3504–3511 (1999).

64. Lin P, M. A., Smith LK, Bigbee C, Bigbee M, Fuhrman C, Grieser H, Chiosea I, Voitenek NN, Capuano SV, Klein E, and L. Flynn J. TNF neutralization results in disseminated disease during acute and latent M. tuberculosis infection with normal granuloma structure. Arthritis Rheumatoid 62, 340–350 (2010).

65. Hawn, T. R., Matheson, A. I., Maley, S. N. & Vandal, O. Host-directed therapeutics for tuberculosis: can we harness the host? Microbiol Mol Biol Rev 77, 608–627 (2013).

66. Wallis, R. S. et al. A study of the safety, immunology, virology, and microbiology of adjunctive etanercept in HIV-1-associated tuberculosis. AIDS 18, 257–264 (2004).

